# Plant lipid metabolism in susceptible and tolerant soybean (*Glycine max*) cultivars in response to *Phytophthora sojae* colonization and infection

**DOI:** 10.1101/2021.06.28.450227

**Authors:** Oludoyin Adeseun Adigun, Thu Huong Pham, Dmitry Grapov, Muhammad Nadeem, Linda Elizabeth Jewell, Mumtaz Cheema, Lakshman Galagedara, Raymond Thomas

**Affiliations:** School of Science and the Environment/ Boreal Ecosystems and Agricultural Sciences, Grenfell Campus, Memorial University of Newfoundland, Corner Brook, Newfoundland and Labrador, A2H 5G4, Canada; CDS-Creative Data Solutions, Colfax, CA, USA; St. John’s Research and Development Centre, Agriculture and Agri-Food Canada, 204 Brookfield Road, St. John’s Newfoundland and Labrador, A1E 6J5, Canada

**Keywords:** *Glycine max* (soybean), membrane lipids, glycerolipids, lipid metabolism, plant-pathogen interaction, *Phytophthora sojae*, root and stem rot, lipid network

## Abstract

Soybean is one of the most cultivated crops globally and a staple food for much of the world’s population. The annual global crop losses due to infection by the *Phytophthora sojae* are currently estimated at approximately $2B USD, yet we have limited understanding of the role of lipid metabolism in the adaptative strategies used to limit infection and crop loss. We employed a multi-modal lipidomics approach to investigate how soybean cultivars remodel their lipid metabolism to successfully limit infection by *Phytophthora sojae*. Both the tolerant and susceptible soybean cultivars showed alterations in lipid metabolism in response to *Phytophthora sojae* infection. Relative to non-inoculated controls, induced accumulation of stigmasterol was observed in the susceptible cultivar whereas, induced accumulation of phospholipids and glycerolipids occurred in tolerant soybean cultivar. We have generated a comprehensive metabolic map of susceptible and tolerant soybean root and stem lipid metabolism to identify lipid modulators of host immune or tolerance response to *Phytophthora sojae* infection and identified potential pathways and unique lipid biomarkers like TG(15:0/22:0/22:5), TG(10:0/10:0/10:0), TG(10:0/10:0/14:0), DG(18:3/18:3), DG(16:0/18:3) and DG(24:0/18:2) as possible targets for the development of future plant protection solutions.

## Introduction

The global population is anticipated to increase to almost 9.7 billion by 2050, which will require a 70% increase in food production (Röös et al. 2017). Food insecurity remains prevalent in many nations despite efforts to improve the production, the quality, and the availability of global food supplies (Matemilola and Elegbede 2017). Food insecurity is a major challenge that must be addressed to meet the demands of an ever-increasing global population (Mc Carthy et al. 2018). To fulfill global food and feed requirements, innovative agricultural practices must be developed to enhance food production, availability and accessibility, which in turn will require advanced knowledge in plant pathology from seedling to crop harvest (Adigun et al. 2020). For instance, plant diseases are caused by infectious pathogens such as fungi, viruses, bacteria, and nematodes (Adigun et al. 2020). These diseases lead to significant annual economic losses in maize, potato, wheat, rice, and soybean worldwide accounting for a 40% yield reduction (Adigun et al. 2020, Fletcher et al. 2006, Savary et al. 2019). Globally, approximately $2B USD are lost annually due to soybean root and stem rot disease caused by the oomycete *Phytophthora sojae* (Dorrance et al. 2003, Ranathunge et al. 2008, Thomas et al. 2007, Wrather et al. 2001). Soybean root and stem rot are the most devastating threat to seedling and plant survival and productivity, particularly during wet growing seasons (Dorrance et al. 2003, Thomas et al. 2007). During the susceptible crop growth stages, pathogens can alter an otherwise favourable environment for the plant into unfavourable conditions, leading to significant yield losses (Velásquez et al. 2018). The repeated applications and heavy dependence on synthetic chemicals such as fungicides limit effective long-term control of this disease, as well as pose serious environmental and human health risks (Dangl et al. 2013). Reducing the frequency and volume of chemical applications in agricultural crops is one of the primary objectives of plant pathological research. Hence, there is a need to develop innovative disease control systems improving the plant’s natural defense mechanisms to build enduring and wide-spectrum disease resistance in crops to improve sustainable agriculture and food security (Brackin et al. 2017, Sui et al. 2008).

Plants respond to different biotic and abiotic stress conditions through various defense mechanisms that may be either constitutive or induced (Adigun et al. 2020, Venegas-Molina et al. 2020). The constitutive system utilizes pre-formed inhibitory chemicals such as alkaloids, saponins, and glycosides, and barriers like wax cuticles, cellulose and suberin to reduce pathogen entry (Adigun et al. 2020, Gao et al. 2014, Thomas et al. 2007). Induced defense mechanisms are triggered by pathogen ingress causing plants to synthesize compounds or enzymes as a result of pathogen detection. This may occur at the site of infection by processes like the oxidative burst or the hypersensitive response, or the production of chitinases, nitric oxide or phytoalexins (Adigun et al. 2020). Furthermore, the response can be systemic in nature, producing pathogenesis-related proteins or the induction of systemic acquired resistance. Plants can also adapt to environmental stresses by regulating biochemical, physiological, and molecular properties of their cellular membrane (Adigun et al. 2020). Several studies have demonstrated the roles of lipids in plant pathology as part of a complex internal defense mechanism in the fight against infections caused by various pathogens (Adigun et al. 2020, Bandara et al. 2019, Lim et al. 2017, Raffaele et al. 2009). Lipid remodeling is a defence mechanism adopted by plants to counteract pathogen attack (Goufo and Cortez 2020). Depending on the composition, lipid molecular species can regulate membrane fluidity, permeability, stability, and integrity during a plant’s response to pathogenic microorganisms. For instance, free fatty acids (FA) such as linoleic acid and oleic acid, which are major components of cellular membranes, play active functions during biosynthesis of the plant cuticular wax, forming the first barrier against pathogens (Lim et al. 2017). Lipid metabolites can also function as intracellular and extracellular signal mediators (Goufo and Cortez 2020, Lim et al. 2017). Plant lipids include glycerophospholipids (GPL), phytosterols (PST), sphingolipids (SGL), glycoglycerolipids (GGL) and glycerolipids (GL) (Adigun et al. 2020, Nadeem et al. 2019, Nadeem et al. 2020), and their metabolites are actively involved in plant defence responses against pathogen colonization (Christensen and Kolomiets 2011, Siebers et al. 2016). They play important roles in the formation of the membrane interface between plant and microbial pathogen (Christensen and Kolomiets 2011, Siebers et al. 2016).

The GPLs of plant membranes possess two FAs as hydrophobic tails at the *sn1* and *sn2* carbons and a hydrophilic head group esterified to a phosphate group at the *sn3* position of the glycerol moiety. The classes of GPLs include phosphatidic acid (PA), phosphatidylcholine (PC), phosphatidylethanolamine (PE), phosphatidylglycerol (PG), phosphatidylinositol (PI), and phosphatidylserine (PS). During plant-pathogen interactions, phospholipid-derived molecules rapidly accumulate and participate in plant signaling and membrane trafficking; they can also activate plant immunity (Canonne et al. 2011, Xing et al. 2020). For instance, PA acts as a novel secondary messenger in plants and its biosynthesis has been reported to be triggered in response to pathogen attack (Laxalt and Munnik 2002, Zhang and Xiao 2015).

Plant sphingolipids are structural components of eukaryotic cellular membranes and play essential roles in maintaining membrane integrity (Ali et al. 2018). They have been recently demonstrated to act as signaling molecules playing crucial functions in the regulation of pathophysiological processes (Berkey et al. 2012, Hannun and Obeid 2018, Heung et al. 2006). Studies have demonstrated that sphingolipids play important roles during biotic stress in plants by activating defences against bacterial and fungal pathogens. For instance, the fungus *Alternaria alternata f. sp. lycopersici* has been shown to activate cell death through disruption of sphingolipid metabolism (Spassieva et al. 2002).

Phytosterols are integral components of cellular membranes and the most abundant sterols in plants include campesterol, sitosterol and stigmasterol (Valitova et al. 2016).Phytosterols are actively involved in regulation of membrane fluidity and integrity, and they influence membrane structural properties and physiological functions of plants. For instance, stigmasterol and beta-sitosterol play a vital role during structural formation and mediate cell membrane functions (Schaller 2004). They have also been demonstrated to play essential roles in plant innate immunity against pathogen attack (Wang et al. 2012).

Galactolipids, including mono-/di-galactosyldiacylglycerol (MGDG and DGDG) are important membrane components in the chloroplasts of eukaryotic plants (Rocha et al. 2018).They play active roles in cell communication, signal transduction, and response to pathogen invasion (Siebers et al. 2016).

Glyceroli pids are actively required during cell growth and cell division (Chapman et al. 2012), serve as energy storage for survival, participate in stress responses, and play an important role in reducing pathogenicity (Murphy 2012). During environmental stresses in plants, triacylglycerol (TG) levels increase as a function of the sequestration of toxic lipid intermediates (Xu and Shanklin 2016). Studies have suggested that diacylglycerols (DGs) serve as signaling molecules during plant growth and development, and in response to stimuli during certain environmental stresses (Dong et al. 2012, Garay et al. 2014). In addition, DG and DG kinase are known to activate immunity during plant defence responses to pathogen attack (Laxalt and Munnik 2002). Although the literature is replete with examples of the plant lipidome mediating plant defence, very little is known concerning how plant lipid metabolism contributes to either successful colonization or tolerance in the soybean-*P. sojae* pathosystem.

We hypothesized that the relative concentrations of membrane lipids in a *P. sojae-*tolerant soybean cultivar would fluctuate more than those of a *P. sojae*-susceptible cultivar following pathogen infection; we hypothesize that these greater changes are just one component of a successful strategy to limit pathogen infection. To this end, we assessed the lipidome of soybean root and stem to understand the functions of lipid metabolism in the response of susceptible and tolerant soybean cultivars to pathogen colonization and infection.

## Results

### Lipid composition of the soybean cultivars in response to *P. sojae* infection

We applied a multi-modal lipidomics approach using UHPLC-C30RP-HESI-HRAM-MS/MS to obtain a detailed understanding of how susceptible and tolerant soybean cultivars remodeled their lipid metabolism to successfully limit infection by *P. sojae* using 10-day old seedlings as a model. The results confirmed as hypothesized that there are significant alterations in the root and stem lipidomes within and between susceptible and tolerant soybean cultivars following inoculation with pathogenic *P. sojae* (Tables 1, 2). Representative chromatograms and mass spectrum demonstrating the separation of the membrane and storage lipids present in the root and stem of the soybean cultivars evaluated (negative and positive ion modes) is shown in Fig. 1. The chromatograms of separated membrane lipids in negative ion mode are shown in Fig.1a. The extracted ion chromatogram of *m/z* 671.46, 802.56 and 833.52 precursor ions of the three selected polar lipids are shown in Fig. 1b. The MS^2^ spectrum of *m/z* 671.46 identified as PA 16:0/18:2 [M-H]^-^ is depicted in Fig. 1c. For example, m/z 152 represent the glycerol moiety (head group) in PA and m/z 255 and 279 represent C16:0 and C18:2 fatty acids present in PA 16:0/18:2 (Fig. 1c). The same convention was used in identifying the other lipids present in Fig 1. This included *m/z* 802.56 identified as PC 16:0/18:2 [M+HCOO]^-^ in Fig. 1d, *m/z* 833.52 representing PI 16:0/18:2 [M-H]^-^ in Fig. 1e. Together, these accounted for some of the main membrane lipids identified in the soybean plant tissue. Similarly, a chromatogram demonstrating the separation of GLs from the stem of the soybean cultivar in the positive ion mode is shown in Fig. 1f. The extracted ion chromatogram of m/z 630.51, 890.72 and 892.74 representing the precursor ions of the three selected GLs are depicted in Fig. 1g. The MS^2^ spectrum of *m/z* 630.51 identified as DG 18:3/18:3 [M+NH_4_]^+^ is depicted in Fig. 1h, the MS^2^ spectrum of *m/z* 802.56 identified as TG 18:3/18:3/18:3 [M+NH_4_]^+^ is depicted in Fig. 1i, and the MS^2^ spectrum of *m/z* 833.52 representing TG 18:3/18:2/18:3 [M+NH_4_]^+^ is depicted in Fig. 1j. These species account for some of the major GLs identified in the plant tissue.

**Table 1.**
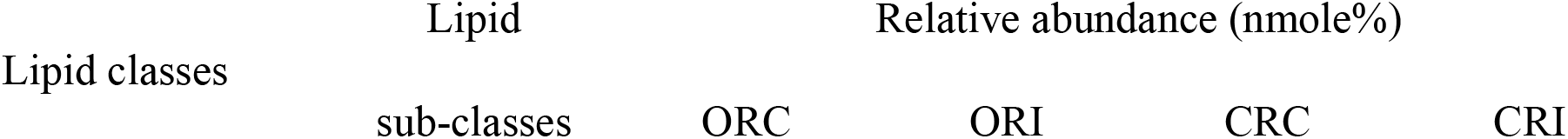
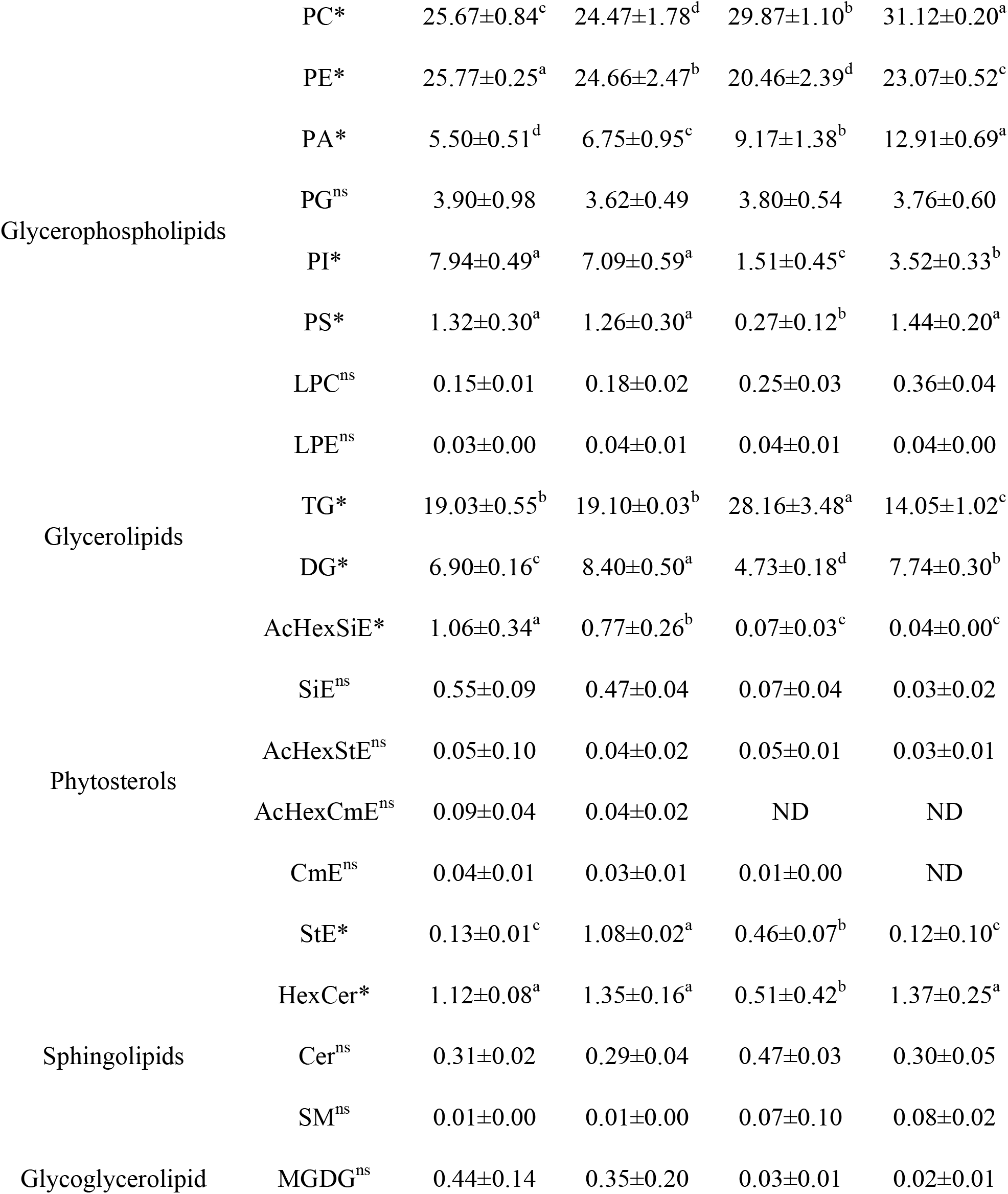
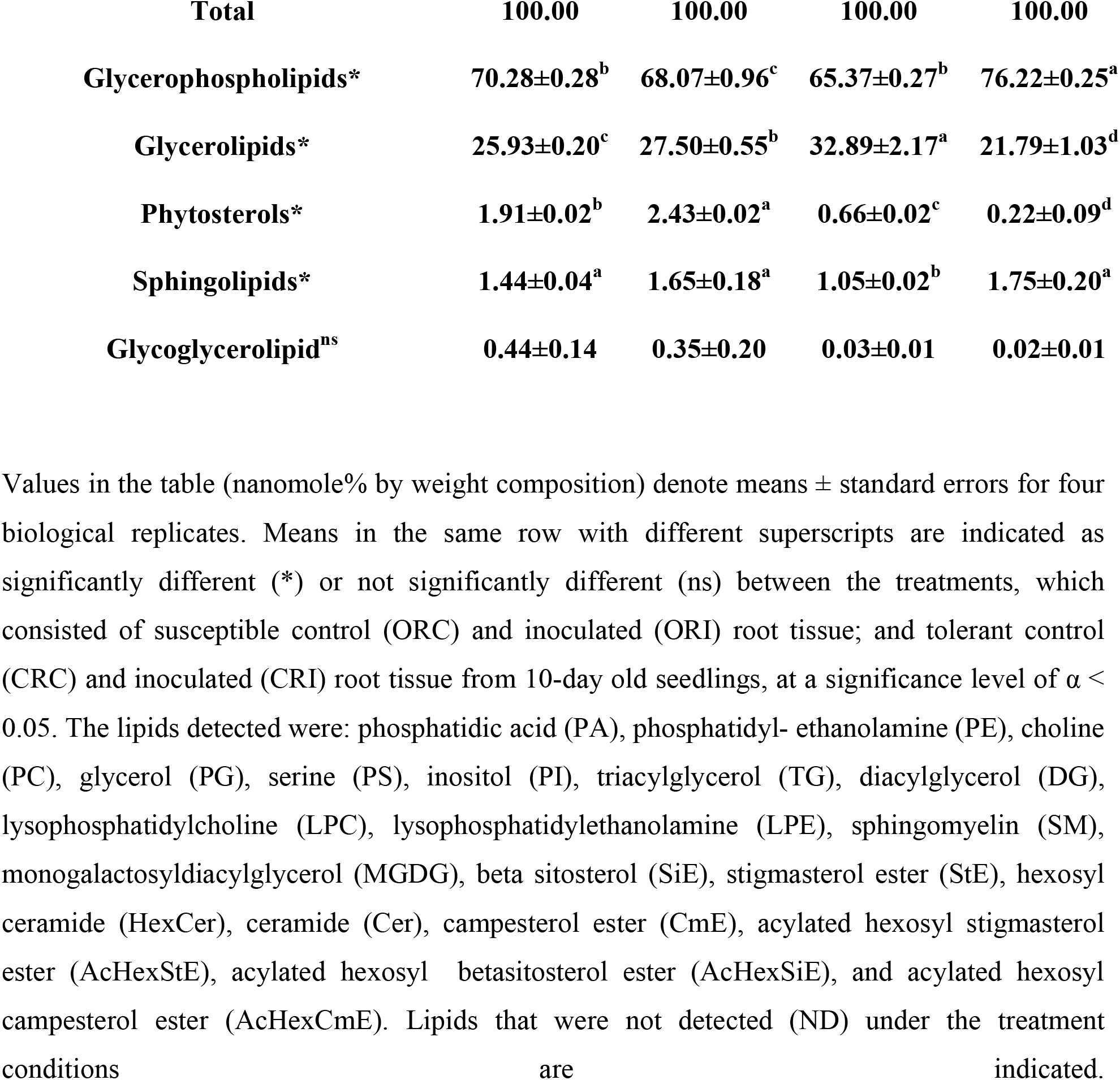
Effect of *Phytophthora sojae* infection on the root lipidome of susceptible (OX760-6) and tolerant (Conrad) soybean cultivars

**Table 2.** Effect of *Phytophthora sojae* infection on stem lipidome of susceptible (OX760-6) and tolerant (Conrad) soybean cultivars

| Lipid classes | Lipid | Relative abundance (nmole%) |  |  |  |
| --- | --- | --- | --- | --- | --- |
|  | sub-classes | OSC | OSI | CSC | CSI |
| Glycerophospholipids | PC* | 22.05±2.50<sup>b</sup> | 17.52±0.92<sup>c</sup> | 20.98±0.26<sup>b</sup> | 23.74±0.39<sup>a</sup> |
|  | PE* | 33.89±2.13<sup>a</sup> | 27.45±1.48<sup>b</sup> | 21.18±0.18<sup>d</sup> | 22.25±0.61<sup>c</sup> |
|  | PA* | 2.72±1.27<sup>d</sup> | 7.60±1.52<sup>c</sup> | 8.39±0.20<sup>b</sup> | 13.37±1.88<sup>a</sup> |
|  | PG* | 8.77±1.16<sup>c</sup> | 7.80±0.41<sup>d</sup> | 12.30±0.69<sup>a</sup> | 10.72±1.56<sup>b</sup> |
|  | PI* | 5.16±0.59<sup>c</sup> | 4.54±0.50<sup>d</sup> | 6.67±0.41<sup>b</sup> | 7.26±0.64<sup>a</sup> |
|  | PS* | 2.24±0.66<sup>b</sup> | 1.25±0.63<sup>c</sup> | 5.33±0.76<sup>a</sup> | 2.91±0.40<sup>b</sup> |
|  | LPC<sup>ns</sup> | 0.42±0.04 | 0.32±0.03 | 0.38±0.04 | 0.36±0.05 |
|  | LPE<sup>ns</sup> | 0.09±0.00 | 0.08±0.02 | 0.09±0.00 | 0.06±0.01 |
| Glycerolipids | TG* | 10.11±1.25<sup>d</sup> | 16.55±0.02<sup>b</sup> | 17.89±1.50<sup>a</sup> | 11.66±2.00<sup>c</sup> |
|  | DG* | 7.78±1.15<sup>b</sup> | 8.35±0.72<sup>a</sup> | 2.62±0.10<sup>d</sup> | 4.45±0.12<sup>c</sup> |
| Phytosterols | AcHexSiE<sup>ns</sup> | 0.55±0.06 | 0.72±0.14 | 0.11±0.02 | 0.60±0.01 |
|  | SiE* | 1.14±0.25<sup>a</sup> | 0.46±0.20<sup>b</sup> | 0.68±0.11<sup>a</sup> | 0.41±0.05<sup>b</sup> |
|  | AcHexStE<sup>ns</sup> | 0.75±0.37 | 0.69±0.21 | 0.04±0.00 | 0.05±0.01 |
|  | AcHexCmE<sup>ns</sup> | 0.14±0.05 | 0.20±0.03 | 0.01±0.00 | ND |
|  | CmE<sup>ns</sup> | 0.44±0.12 | 0.15±0.05 | 0.01±0.00 | 0.01±0.01 |
|  | StE* | 0.34±0.16<sup>b</sup> | 1.98±0.02<sup>a</sup> | 0.33±0.05<sup>b</sup> | 0.40±0.15<sup>b</sup> |
| Sphingolipids | HexCer* | 0.89±0.17<sup>b</sup> | 1.96±0.02<sup>a</sup> | 1.00±0.08<sup>b</sup> | 1.67±0.03<sup>a</sup> |
|  | Cer<sup>ns</sup> | 0.27±0.06 | 0.31±0.02 | 0.71±0.04 | 0.21±0.03 |
|  | SM<sup>ns</sup> | ND | ND | 0.03±0.01 | 0.01±0.00 |
| Glyceroglycolipid | MGDG* | 2.25±0.08<sup>a</sup> | 2.09±0.06<sup>a</sup> | 1.25±0.18<sup>b</sup> | 0.47±0.07<sup>c</sup> |
| **Total** |  | **100.00** | **100.00** | **100.00** | **100.00** |
| **Glycerophospholipids\*** |  | **75.34±1.20<sup>b</sup>** | **66.56±1.32<sup>c</sup>** | **75.32±1.22<sup>b</sup>** | **80.67±2.15<sup>a</sup>** |
| **Glycerolipids\*** |  | **17.89±0.25<sup>c</sup>** | **24.90±1.51<sup>a</sup>** | **20.51±1.60<sup>b</sup>** | **16.11±1.13<sup>d</sup>** |
| **Phytosterols\*** |  | **3.36±0.12<sup>b</sup>** | **4.18±0.66<sup>a</sup>** | **1.18±0.19<sup>c</sup>** | **0.86±0.06<sup>d</sup>** |
| **Sphingolipids\*** |  | **1.16±0.03<sup>c</sup>** | **2.27±0.05<sup>a</sup>** | **1.74±0.12<sup>b</sup>** | **1.89±0.06<sup>b</sup>** |
| **Glyceroglycolipid\*** |  | **2.25±0.08<sup>a</sup>** | **2.09±0.06<sup>a</sup>** | **1.25±0.18<sup>b</sup>** | **0.47±0.07<sup>c</sup>** |

Values in the table (nanomole% by weight composition) denote means ± standard errors for four
biological replicates. Means in the same row with different superscripts are indicated as
significantly different (\*) or not significantly different (ns) between the treatments, which
consisted of susceptible control (OSC) and inoculated (OSI) stem tissue; and tolerant control
(CSC) and inoculated (CSI) stem tissue from 10-day old seedlings, at a significance level of $\alpha <$
0.05. The lipids detected were: phosphatidic acid (PA), phosphatidyl- ethanolamine (PE), choline
(PC), glycerol (PG), serine (PS), inositol (PI), triacylglycerol (TG), diacylglycerol (DG),
lysophosphatidylcholine (LPC), lysophosphatidylethanolamine (LPE), sphingomyelin (SM),
monogalactosyldiacylglycerol (MGDG), beta sitosterol (SiE), stigmasterol ester (StE), hexosyl

**Fig. 1.**
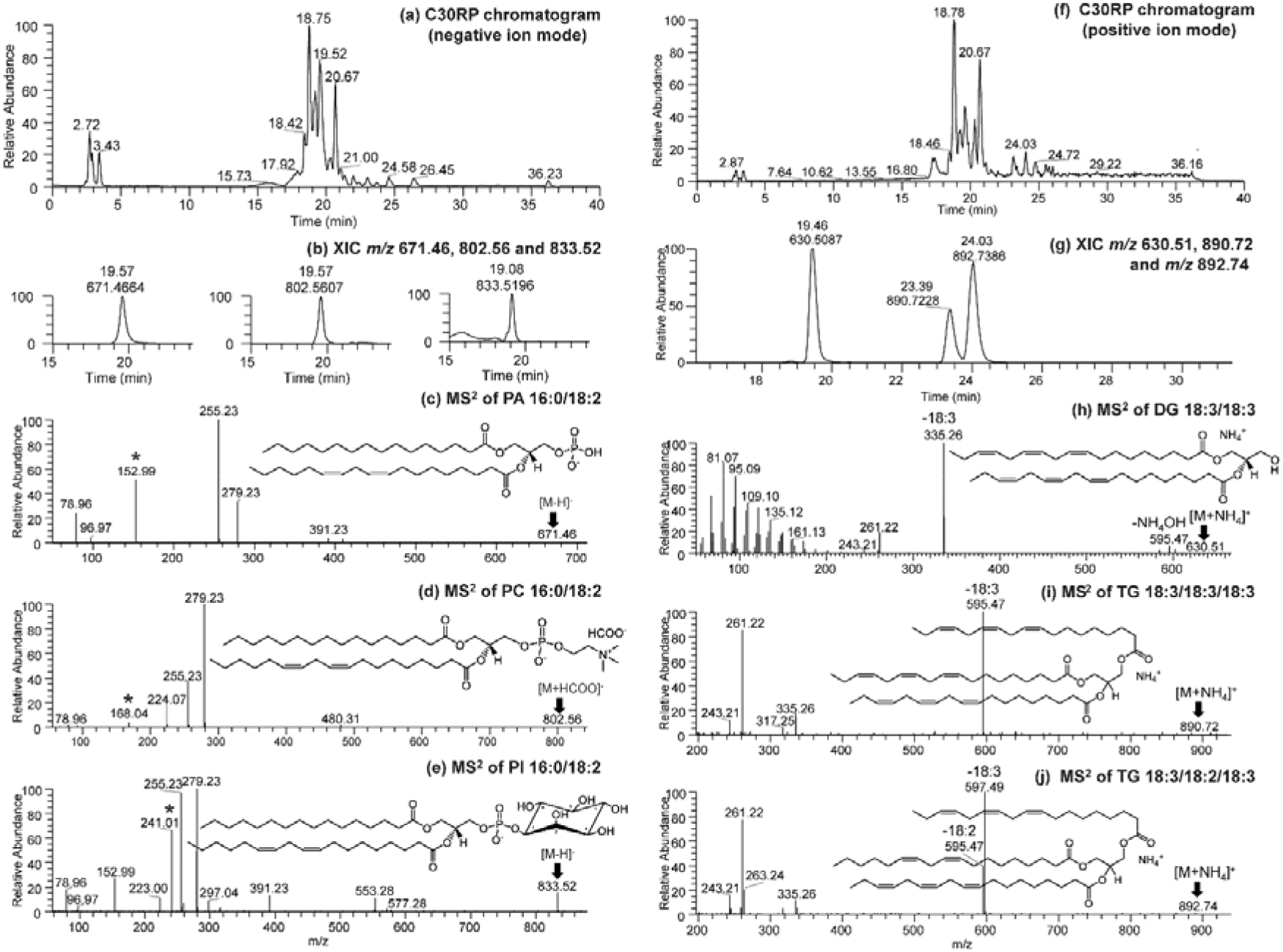
Chromatogram demonstrating the UHPLC-C30RP-HESI-HRAM-MS separation of the membrane lipids and glycerolipids in the root and stem of susceptible and tolerant soybean cultivars. (a) LC-MS chromatogram of separated membrane lipids in negative ion mode, (b) Extracted ion chromatogram (XIC) of *m/z* 671.46, 802.56 and 833.52 precursor ions of the three selected polar lipids, (c) MS^2^ spectrum of *m/z* 671.46 identified as PA 16:0/18:2 [M-H]^-^, (d) MS^2^ spectrum of *m/z* 802.56 identified as PC 16:0/18:2 [M+HCOO]^-^ and (e) MS^2^ spectrum of *m/z* 833.52 representing PI 16:0/18:2 [M-H]^-^ identified in the negative ion mode; (f) LC-MS chromatogram in positive ion mode of separated glycerolipids in positive ion mode (g) Extracted ion chromatogram (XIC) of *m/z* 630.51, 890.72 and 892.74 precursor ions of the three selected glycerolipids, (h) MS^2^ spectrum of *m/z* 630.51 identified as DG 18:3/18:3 [M+NH_4_]^+^, (i) MS^2^ spectrum of *m/z* 802.56 identified as TG 18:3/18:3/18:3 [M+NH_4_]^+^ and (j) MS^2^ spectrum of *m/z* 833.52 representing TG 18:3/18:2/18:3 [M+NH_4_]^+^ identified in the positive ion mode. PA = phosphatidic acid, PC = phosphatidylcholine, and PI = phosphatidylinositol, DG = diacylglycerol, TG = triacylglycerol, and * represent the head group for each of the lipid class presented.

We observed five lipid classes: GPL, PST, GL, SGL, and GGL in soybean stem and root. Glycerophospholipids accounted for the highest portion of total lipids in both cultivars, irrespective of tissue type or inoculation status, representing 65.37±0.27 nmol% to 76.22±0.25 nmol% of all lipids in root (Table 1) and 66.56±1.32 to 80.67±2.15 nmol% in stem (Table 2), followed by GLs which ranged from 21.79±1.03 nmol% to 32.89±2.17 nmol% in the roots and 16.11±1.13 nmol% to 24.90±1.51 nmol% in the stems (Table 2). Phytosterols, SGLs, and GGLs were present in lower quantities ranging between 0.02±0.01 nmol% to 2.43±0.02 nmol% for root (Table 1) and 0.47±0.07 nmol% to 4.18±0.66 nmol% for stem (Table 2). From the five lipid classes investigated, 20 subclasses were analyzed across both root and stem which include eight GPLs, two GLs, six PSTs, three SGLs, and one GGL (Tables 1, 2). In tolerant root tissue, the percentage of the following lipids increased after inoculation: PC (4.18%), PE (12.76%), PA (40.79%), PI (133.11%), PS (433.33%), hexaceramide (HexCer; 168.63%), and DG (63.64%) (Table 1). In contrast, the following lipid increases were observed in the susceptible roots: PA (22.73%), DG (21.74%) and stigmasterol ester (StE; 730.77%) (Table 1). In the stem of the tolerant cultivar, an increase in lipid levels was observed for PC (13.16%), PE (5.05%), PA (59.36%), PI (8.85%), HexCer (67.00%), and DG (69.85%) whereas in susceptible cultivar’s stem, an increase in lipid levels was observed for PA (179.41%), DG (7.33%), TG (63.70%), HexCer (120.22%) and StE (482.35%) (Table 2). Specifically, we observed significantly higher levels of major GPLs, HexCer and DG in the tolerant cultivar, but higher levels of TG and StE in the susceptible cultivar in response to *P. sojae* colonization and infection.

### Modification of membrane lipids in soybean cultivars in response to *P. sojae* infection

An analysis of membrane lipids in soybean root and stem tissues following infection with *P. sojae* was performed to determine changes and modification of membrane lipids during host-pathogen interaction. Figs. 2a-d and 3a-d demonstrate the changes that occurred in membrane lipids during host-pathogen interactions. Based upon the membrane lipid molecular species observed, we conducted PLS-DA to determine the most important membrane lipid molecular species with influential loadings (Figs. 2a, b and 3a, b) segregating the tolerant from the susceptible cultivar based on pathogen challenge. The model quality (Q^2^) represents 95% and 96% variability in root and stem, respectively (Fig. 2a, 3a). The result from the PLS-DA observation plot showed the segregation of the susceptible and tolerant soybean cultivars before and after infection into four distinct groups that are in accordance with the root and stem membrane lipid molecular species (Fig. 2b, 3b). The root membrane lipid molecular species (Fig. 2b) separated the treatments into four distinct quadrants (Q). Quadrant 1 contained the lipid molecular species associated with Conrad root control (CRC) treatment, Q-2 contained Conrad root inoculated (CRI) treatment, Q-3 contained OX760-6 root control (ORC) and Q-4 had the OX760-6 root inoculated (ORI) treatment, respectively. Similarly, the changes in soybean stem (Fig. 3b), lipid molecular species separated the treatments into 4 distinct quadrants (Q-1, Q-2, Q-3 and Q-4) consisting of Conrad stem control (CSC), Conrad stem inoculated (CSI), OX760-6 stem control (OSC) and OX760-6 stem inoculated (OSI) treatments, respectively.

**Fig. 2.**
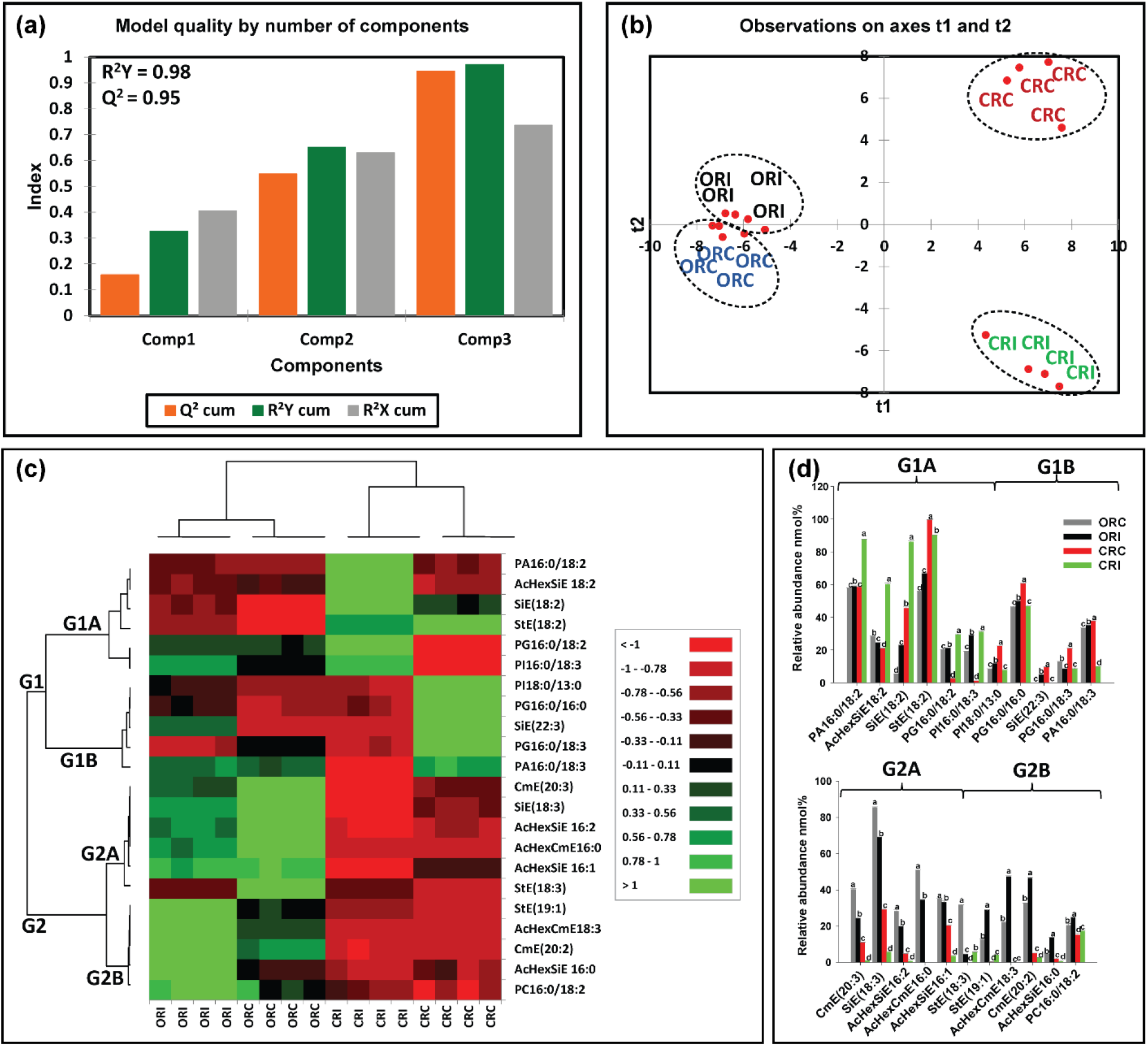
Differences in root membrane lipids in susceptible (OX760-6) and resistant (Conrad) soybean cultivars inoculated with *P. sojae* relative to control plants. (**a**) Model quality for partial least squares-discriminant analysis (PLS-DA); (**b**) Observation plot based upon differences in molecular species in root membrane lipids of OX760-6 and Conrad cultivars; (**c**) Heat map demonstrating clusters of root membrane lipid species in OX760-6 and Conrad cultivars treated or untreated with *P. sojae*. Each cultivar and treatment were grouped separately using ascendant hierarchical cluster analysis based upon Euclidian distance at interquartile range of 0.15. The left columns denote the cluster segregated root membrane lipid species, while the above columns segregated soybean cultivars based upon similarities in abundance. The abundance of root membrane lipid species is denoted using color: red for lower level, black for intermediate level, and green for higher level. Group 1 and 2 (G1 and G2) and subgroups (G1A, G1B, G2A and G2B) are root membrane lipid species that were accountable for the formation of clustered patterns in the heat map that were applied for determination of significant differences between the soybean cultivars (OX760-6 and Conrad) root membrane lipid species in each of the bar chart (Fig. 1d) beside the heat map; and (**d**) Bar charts describe the relative abundance of root membrane lipid species as a mean nmol% ± SE (n = 4). Significant differences between root membrane lipid species are indicate using letter a-d on top of the bars as described by Fisher’s LSD multiple comparisons test using ANOVA (α = 0.05). The G1 and G2, and G1A, G1B, G2A and G2B are root membrane lipid species that were accountable for the formation of clustered patterns in the heat map that were applied for the determination of significant differences between the soybean cultivars (OX760-6 and Conrad) root membrane lipid species as illustrated in the bar charts.

**Fig. 3.**
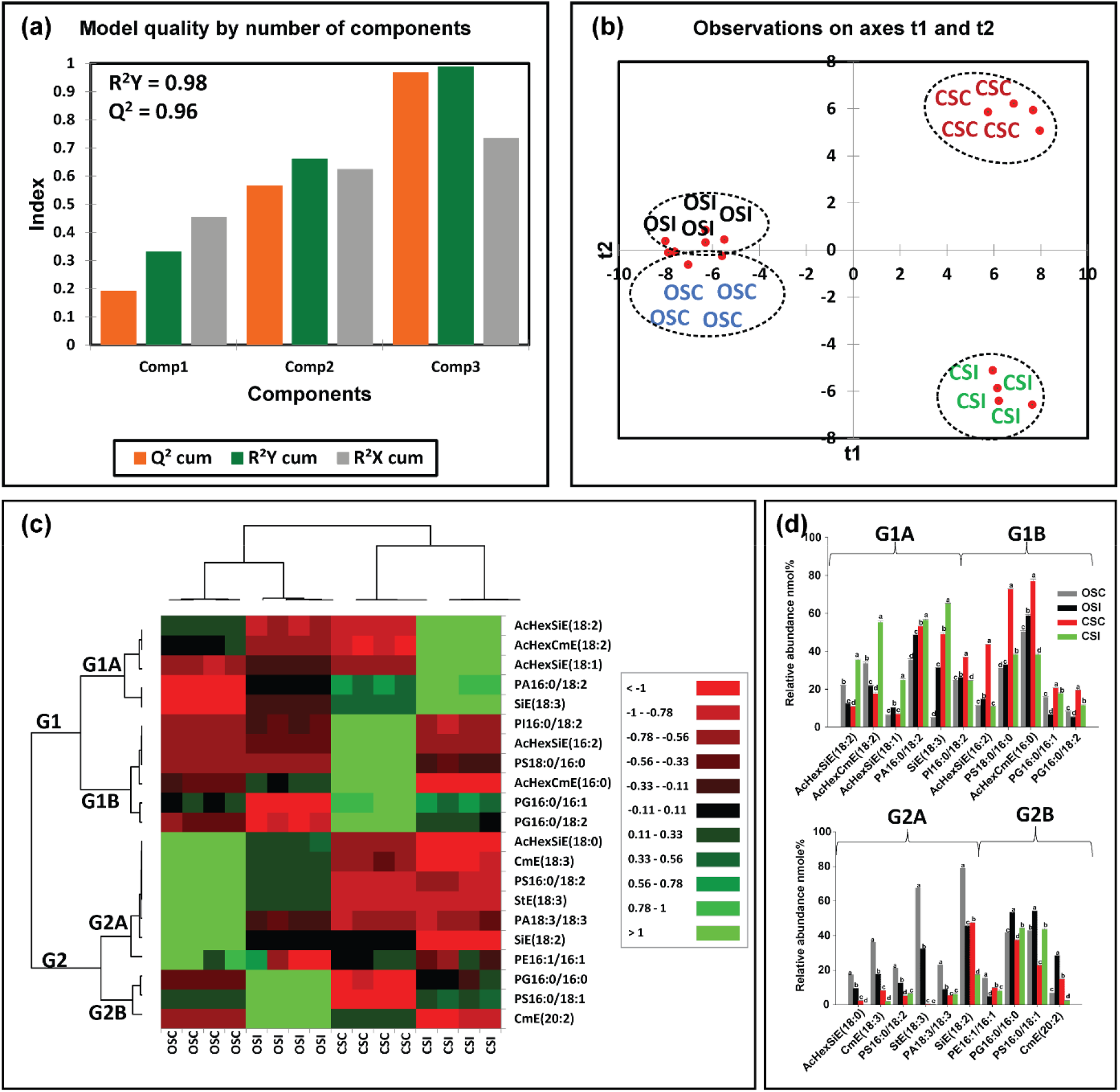
Differences in stem membrane lipids in susceptible (OX760-6) and resistant (Conrad) soybean cultivars inoculated with *P. sojae* relative to control plants. (**a**) Model quality for partial least squares-discriminant analysis (PLS-DA); (**b**) Observation plot based upon differences in molecular species in stem membrane lipids of OX760-6 and Conrad cultivars; (**c**) Heat map demonstrating clusters of stem membrane lipid species in OX760-6 and Conrad cultivars treated or untreated with *P. sojae*. Each cultivar and treatment were grouped separately using ascendant hierarchical cluster analysis based upon Euclidian distance at interquartile range of 0.15. The left columns denote the cluster segregated stem membrane lipid species, while the above columns segregated soybean cultivars based upon similarities in abundance. The abundance of stem membrane lipid species is denoted using color: red for lower level, black for intermediate level, and green for higher level. Group 1 and 2 (G1 and G2) and subgroups (G1A, G1B, G2A and G2B) are stem membrane lipid species that were accountable for the formation of clustered patterns in the heat map that were applied for determination of significant differences between the soybean cultivars (OX760-6 and Conrad) stem membrane lipid species in each of the bar chart (Fig. 1d) beside the heat map; and (**d**) Bar charts describe the relative abundance of stem membrane lipid species as a mean nmol% ± SE (n = 4). Significant differences between stem membrane lipid species are indicate using letter a-d on top of the bars as described by Fisher’s LSD multiple comparisons test using ANOVA (α = 0.05). The G1 and G2, and G1A, G1B, G2A and G2B are stem membrane lipid species that were accountable for the formation of clustered patterns in the heat map that were applied for the determination of significant differences between the soybean cultivars (OX760-6 and Conrad) stem membrane lipid species as illustrated in the bar charts.

Based upon Component 3, which demonstrated the highest variation in the data (Figs. 2a, 3a), 22 membrane lipid molecular species from root tissue and 21 membrane lipid molecular species from stem tissue were selected for further analysis. Heat maps (Figs. 2c, 3c) were generated for the lipids with influential loadings accounting for the genotype and treatment segregation to further classify the treatments based on the altered membrane lipidome following infection. The cut-off value for variables important in projection (VIP) scores was defined as >1 (Nadeem et al. 2020, Ravipati et al. 2015). The 22 important root membrane lipid molecular species and 21 important stem membrane lipid molecular species were selected based on VIP scores greater than 1. The output from the heat map analysis showed four different clusters of the soybean root and stem membrane lipid molecular species following inoculation with *P. sojae* (Figs. 2c, 3c).

The heat map clusters root membrane lipid species into two main groups (G), G1 and G2, and four sub-groups, G1A, G1B, G2A and G2B. These groupings distinguished the susceptible cultivar (ORC & ORI) from the tolerant cultivar (CRC & CRI) in the root membrane lipid species in response to infection. We observed differences in the root membrane lipid species in G1A, where the relative abundance (nmol%) of PA(16:0/18:2), AcHexSiE(18:2), SiE(18:2), PG(16:0/18:2), PI(16:0/18:3) were significantly elevated in the tolerant cultivar challenged with *P. sojae* relative to the control and the susceptible cultivar (Fig. 2c). Lipid molecular species belonging to group G1B {PI(18:0/13:0), PG(16:0/16:0), SiE(22:3), PG(16:0/18:3), and PA(16:0/18:3)} were significantly lower in the tolerant cultivar that was challenged with the pathogen, whereas there was no difference in the susceptible cultivar whether treated or untreated with the pathogen (Fig. 2c). Lipid molecular species belonging to group G2A {CmE(20:3), SiE(18:3), AcHexSiE(16:2), AcHexCmE(16:0), AcHexSiE(16:1) and StE(18:3)} were not different in the root of the tolerant cultivar when treated or untreated with the pathogen, but were significantly lower in the root of susceptible cultivar challenged with the pathogen (Fig. 2c). Finally, in G2B, the relative abundances of StE(19:1), AcHexCmE(18:3), CmE(20:2), AcHexSiE(16:0), and PC(16:0/18:2) were not significantly different in the root of the tolerant cultivar but were significantly higher in the root of the susceptible cultivar infected by the pathogen (Fig. 2c). These data are corroborated by Fig. 2d, which demonstrates the significant differences in the molecular species in the root of tolerant and susceptible cultivars. In the pathogen challenged roots of the tolerant cultivar, the relative abundances of PA(16:0/18:2), AcHexSiE(18:2), PG(16:0/18:2), PG(16:0/18:3), (StE18:3) and (PC(16:0/18:2) were higher, whereas the relative abundances of StE (18:2), SiE(22:3), StE (19:1), AcHexCmE(18:3), CmE(20:2), AcHexSiE(16:0), and (PC(16:0/18:2) were lower in the root of susceptible cultivar infected with the pathogen (Fig. 2d).

Similarly, the heat map clusters stem membrane lipid molecular species into two major groups (G1and G2) which are further divided into sub-groups G1A, G1B, G2A and G2B. These groupings distinguished the susceptible cultivar (OSC & OSI) from the tolerant cultivar (CSC & CSI) in the stem membrane lipid molecular species. We observed stem membrane lipid molecular species in the tolerant cultivar, corresponding to G1A and consisting of AcHexSiE(18:2), AcHexCmE(18:2), AcHexSiE(18:1), PA(16:0/18:2), and SiE(18:3), were significantly higher in the tolerant cultivar challenged with *P. sojae* but there were no significant differences in the lipids of the susceptible cultivar. On the other hand, PA(16:0/18:2) was higher in the pathogen-infected tissue relative to the control (Fig. 3c). Lipid molecular species belonging to G1B {(PI(16:0/18:2), AcHexSiE(16:2), PS(18:0/16:0), AcHexCmE(16:0), PG(16:0/16:1), and PG(16:0/18:2)} were significantly lower in the tolerant cultivar challenged with the pathogen, whereas there was no difference in the susceptible cultivar regardless of infection status (Fig. 3c). Lipid molecular species belonging to G2A {AcHexSiE(18:0), CmE(18:3), PS(16:0/18:2), StE(18:3), PA(18:3/18:3), SiE(18:2) and PE(16:1/16:1)} were not significantly different in the stem of the tolerant cultivar but were significantly lower in the stem of susceptible cultivar challenged with the pathogen. Finally, in G2B, the levels of PG(16:0/16:0), PS(16:0/18:1) and CmE(20:2) significantly increased in the stem of the susceptible cultivar challenged with *P. sojae* (Fig. 3c). These trends are further corroborated by the output presented in Fig. 3d, which demonstrates the significant differences in the molecular species in the stem of tolerant and susceptible cultivar when challenged with the pathogen. For example, AcHexSiE(18:2), AcHexCmE(18:2), AcHexSiE(18:1), SiE(18:3), PS(16:0/18:2), and PA(18:3/18:3) were significantly higher in the stem of the tolerant cultivar, whereas AcHexSiE(18:1), AcHexSiE(16:2), AcHexCmE(16:0), and CmE(20:2) were significantly higher in the stem of the susceptible cultivar (Fig. 3d). These results showed there were significantly higher levels of GPL molecular species in root and stem of tolerant cultivar whereas there were significantly higher relative levels of PST molecular species in the root and stem of the susceptible cultivar in response to infection by the pathogen.

### Modification of glycerolipids in soybean cultivars in response to *P. sojae* infection

We also analysed GL in soybean root and stem tissues following infection with *P. sojae* to determine whether their levels and composition were altered during host-pathogen interaction (Figs. 4a-d, 5a-d). Triacylglycerols and DGs were observed to be the major GLs present regardless of soybean cultivar. We next performed PLS-DA to identify the most important TG and DG species with influential loadings (Figs. 4a, 4b, 5a, 5b) segregating the tolerant and susceptible soybean cultivars in their response to *P. sojae* colonization and infection. The model quality (Q^2^) represents 80% and 83% variability in root and stem, respectively (Fig. 4a, 5a). The result from the PLS-DA observation plot showed the segregation of the susceptible and tolerant soybean cultivars that were infected or not infected with the pathogen into four distinct quadrants based on the levels of GL molecular species (Figs. 4b, 5b). The root GL molecular species (Fig. 3b) separated the treatments into four distinct quadrants. Quadrants 1-4 were composed of the GL molecular species of CRC, CRI, ORC and ORI treatments, respectively. Similar to the changes in soybean stem (Fig. 5b), GL species separated the treatments into 4 distinct quadrants (Q1-Q4) consisting of the GLs from CSC, CSI, OSC and OSI, respectively.

**Fig. 4.**
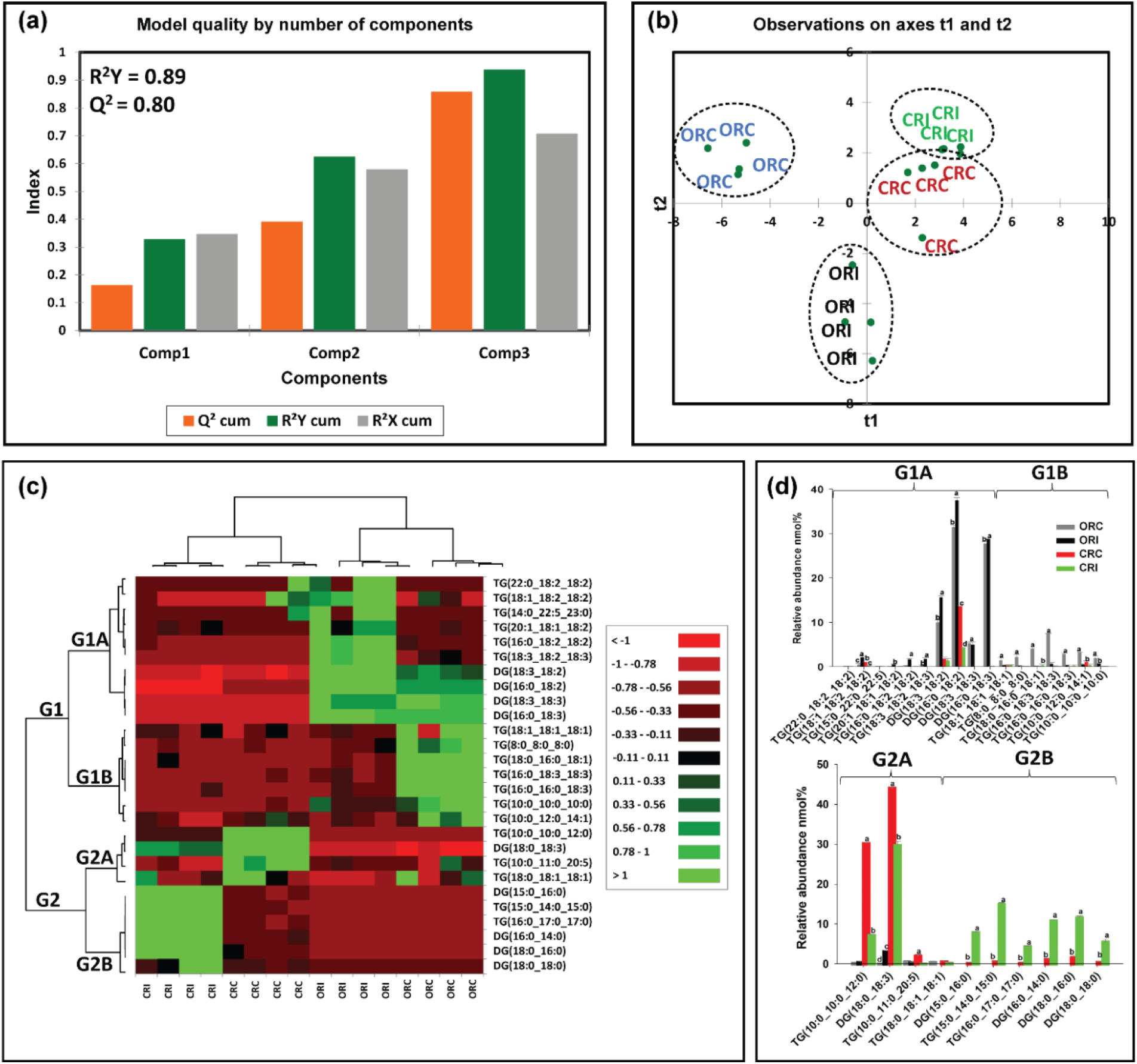
Differences in root glycerolipid species in susceptible (OX760-6) and resistant (Conrad) soybean cultivars inoculated with *P. sojae* relative to control plants. (**a**) Model quality for partial least squares-discriminant analysis (PLS-DA); (**b**) Observation plot based upon differences in molecular species in root glycerolipid species of OX760-6 and Conrad cultivars; (**c**) Heat map demonstrating clusters of root glycerolipid species in OX760-6 and Conrad cultivars treated or untreated with *P. sojae*. Each cultivar and treatment were grouped separately using ascendant hierarchical cluster analysis based upon Euclidian distance at interquartile range of 0.15. The left columns denote the cluster segregated root glycerolipid species, while the above columns segregated soybean cultivars based upon similarities in abundance. The abundance of root glycerolipid species is denoted using color: red for lower level, black for intermediate level, and green for higher level. Group 1 and 2 (G1 and G2) and subgroups (G1A, G1B, G2A and G2B) are root glycerolipid species that were accountable for the formation of clustered patterns in the heat map that were applied for determination of significant differences between the soybean cultivars (OX760-6 and Conrad) root glycerolipid species in each of the bar chart (Fig. 1d) beside the heat map; and (**d**) Bar charts describe the relative abundance of root glycerolipid species as a mean nmol% ± SE (n = 4). Significant differences between root glycerolipid species are indicate using letter a-d on top of the bars as described by Fisher’s LSD multiple comparisons test using ANOVA (α = 0.05). The G1 and G2, and G1A, G1B, G2A and G2B are root glycerolipid species that were accountable for the formation of clustered patterns in the heat map that were applied for the determination of significant differences between the soybean cultivars (OX760-6 and Conrad) root glycerolipid species as illustrated in the bar charts.

**Fig. 5.**
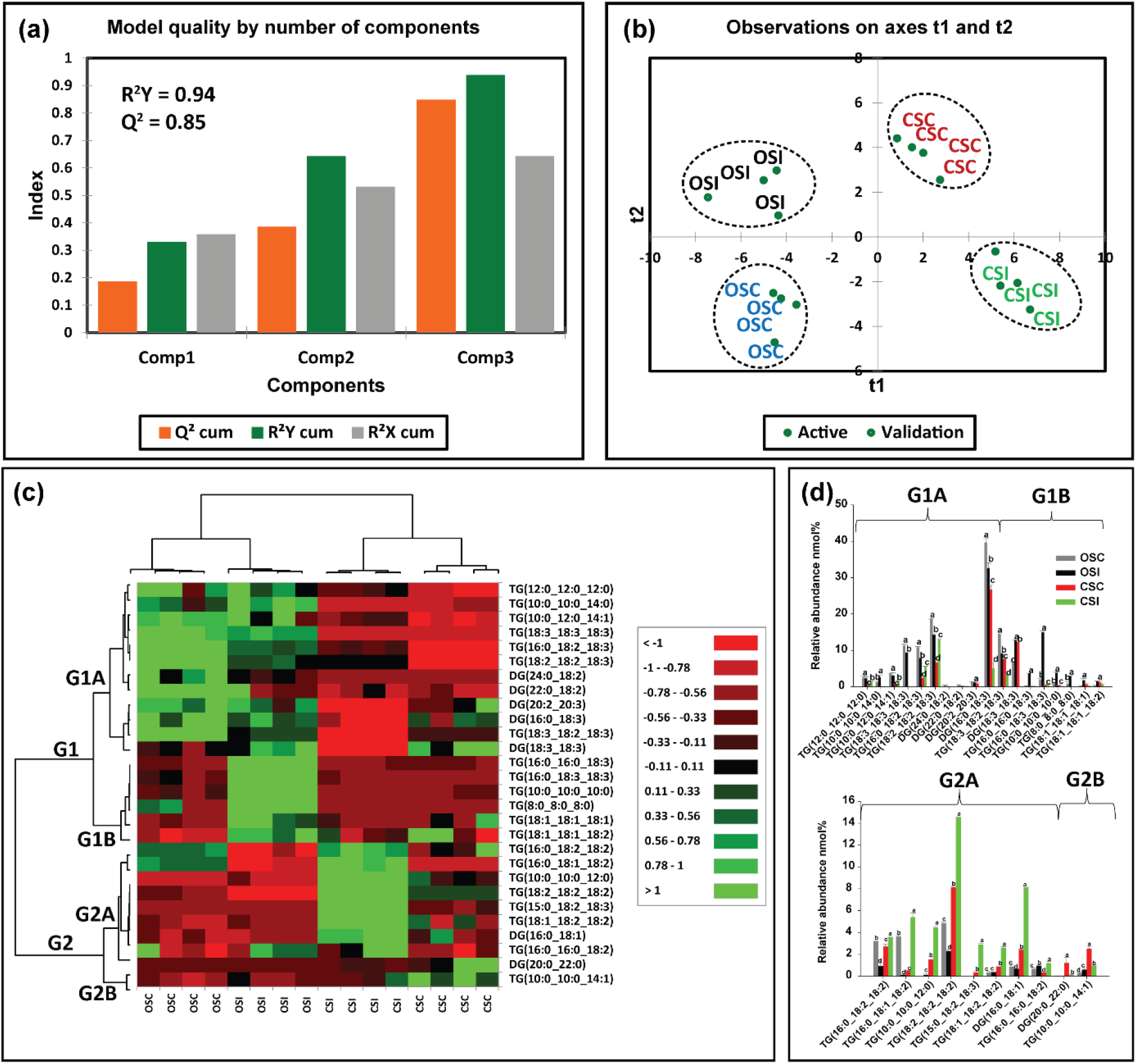
Differences in stem glycerolipid species in susceptible (OX760-6) and resistant (Conrad) soybean cultivars inoculated with *P. sojae* relative to control plants. (**a**) Model quality for partial least squares-discriminant analysis (PLS-DA); (**b**) Observation plot based upon differences in molecular species in stem glycerolipid species of OX760-6 and Conrad cultivars; (**c**) Heat map demonstrating clusters of stem glycerolipid species in OX760-6 and Conrad cultivars treated or untreated with *P. sojae*. Each cultivar and treatment were grouped separately using ascendant hierarchical cluster analysis based upon Euclidian distance at interquartile range of 0.15. The left columns denote the cluster segregated stem glycerolipid species, while the above columns segregated soybean cultivars based upon similarities in abundance. The abundance of stem glycerolipid species is denoted using color: red for lower level, black for intermediate level, and green for higher level. Group 1 and 2 (G1 and G2) and subgroups (G1A, G1B, G2A and G2B) are stem glycerolipid species that were accountable for the formation of clustered patterns in the heat map that were applied for determination of significant differences between the soybean cultivars (OX760-6 and Conrad) stem glycerolipid species in each of the bar chart (Fig. 1d) beside the heat map; and (**d**) Bar charts describe the relative abundance of stem glycerolipid species as a mean nmol% ± SE (n = 4). Significant differences between stem glycerolipid species are indicate using letter a-d on top of the bars as described by Fisher’s LSD multiple comparisons test using ANOVA (α = 0.05). The G1 and G2, and G1A, G1B, G2A and G2B are stem glycerolipid species that were accountable for the formation of clustered patterns in the heat map that were applied for the determination of significant differences between the soybean cultivars (OX760-6 and Conrad) stem glycerolipid species as illustrated in the bar charts.

Based upon component 3 which explained the highest level of variation in the data (Figs. 4a, 5a), 27 GL molecular species from root tissues and 28 GL molecular species from the stem tissue with VIPs greater than 1 were selected for further multivariate analysis. Heat maps (Figs. 4c, 5c) were next generated for the lipids with influential loadings accounting for the genotype and treatment segregation to further classify the treatments based on the altered GL in the infected tissue. The output from the heat map analysis showed four different clusters of the soybean root and stem membrane lipid molecular species following inoculation with *P. sojae* (Figs. 4c, 5c). The heat map clustered GL species into two main groups, G1 and G2, and four sub-groups (G1A, G1B, G2A and G2B). These groupings distinguished the GL lipid molecular species in the root of the susceptible cultivar (ORC and ORI) from those of the root of the tolerant cultivar (CRC and CRI), as well as the stem-derived GL lipid molecular species from both the susceptible (OSC and OSI) and tolerant cultivar (CSC and CSI) (Figs 4-5).

We observed that root GL molecular species in G1A {TG(22:0/18:2/18:2), TG(18:1/18:2/18:2), TG(14:0/22:5/23:0), TG(20:1/18:1/18:2), TG(16:0/18:2/18:2), TG(18:3/18:2/18:3), DG(18:3/18:2), DG(16:0/18:2), DG(18:3/18:3), and DG(16:0/18:3)} did not differ in the tolerant cultivar challenged with *P. sojae* relative to control, but were significantly higher in the susceptible cultivar challenged with the pathogen (Fig. 4c). Lipid molecular species belonging to group G1B {TG(18:1/18:1/18:1), TG(8:0/8:0/8:0), TG(18:0/16:0/18:1), TG(16:0/18:3/18:3), TG(16:0/16:0/18:3), TG(10:0/12:0/14:1), and TG(10:0/10:0/10:0)} also did not differ in the tolerant cultivar regardless of infection status, but were significantly lower in the susceptible cultivar in response to infection (Fig. 4c). In contrast, lipid molecular species belonging to group G2A {TG(10:0/10:0/12:0), DG(18:0/18:3), TG(10:0/11:0/20:5), and TG(18:0/18:1/18:1)} were significantly lower in the root of the tolerant cultivar that was challenged with the pathogen, but no differences were observed for the susceptible cultivar regardless of infection status (Fig. 4c). Finally, in G2B, the relative abundances of DG(15:0/16:0), TG(15:0/14:0/15:0), TG(16:0/17:0/17:0), DG(16:0/14:0), DG(18:0/16:0) and DG(18:0/18:0) were significantly higher in the tolerant cultivar in response to infection, whereas no differences were observed for the susceptible cultivar regardless of infection status(Fig. 4c). These data are corroborated by Fig. 4d, which demonstrates the significant differences in the molecular species in the root of tolerant and susceptible cultivars. In response to pathogen challenge, TG(18:0/16:0/18:1), DG(15:0/16:0), TG(15:0/14:0/15:0), TG(16:0/17:0/17:0), DG(16:0/14:0), DG(18:0/16:0) and DG(18:0/18:0) were significantly higher in the root of the tolerant cultivar while TG(18:1/18:2/18:2), TG(20:1/18:1/18:2), TG(16:0/18:2/18:2), TG(18:3/18:2/18:3), DG(18:3/18:2), DG(16:0/18:2), DG(18:0/18:3) were significantly higher in the root of the susceptible cultivar after infection (Fig. 4d).

Likewise, the heat map clusters stem GL lipid molecular species into G1and G2, and sub-groups G1A, G1B, G2A and G2B. These groupings distinguished the susceptible cultivar from the tolerant cultivar in the stem GL molecular species. We observed stem GL lipid molecular species that belonged to G1A {TG(12:0/12:0/12:0), TG(10:0/10:0/14:0), TG(10:0/10:0/14:1), TG(18:3/18:3/18:3), TG(16:0/18:2/18:3), TG(18:2/18:2/18:3), DG(24:0/18:2), DG(22:0/18:2), DG(20:2/20:3), DG(16:0/18:3) TG(18:3/18:2/18:3), and DG(18:3/18:3)} did not change in the tolerant cultivar challenged with *P. sojae* relative to the control, but were significantly lower in the susceptible cultivar that had been infected (Fig. 5c). Lipid molecular species belonging to group G1B {TG(16:0/16:0/18:3), TG(16:0/18:3/18:3), TG(10:0/10:0/10:0), TG(8:0/8:0/8:0), TG(18:1/18:1/18:1), TG(18:1/18:1/18:2)} also did not differ among the tolerant cultivar, but were significantly higher in the susceptible cultivar that had been treated with the pathogen (Fig. 5c). In contrast, lipid molecular species belonging to group G2A {TG(16:0/18:2/18:2), TG(16:0/18:1/18:2), TG(10:0/10:0/12:0), TG(18:2/18:2/18:2), TG(15:0/18:2/18:3), TG(18:1/18:2/18:2), DG(16:0/18:1), and TG(16:0/16:0/18:2)} were significantly higher in the stem of the tolerant cultivar that had been challenged with the pathogen, but no significant differences were observed in the stem of susceptible cultivar (Fig. 5c). Finally, in G2B, the relative abundances of DG(20:0/22:0) and TG(10:0/10:0/14:1) were significantly lower in the stem of the tolerant cultivar when challenged with *P. sojae* but did not differ among the susceptible cultivar (Fig. 5c). These data are corroborated by Fig. 5d, which demonstrates the significant differences in the GL molecular species in the stem of tolerant and susceptible cultivars. In response to pathogen challenge, TG(12:0/12:0/12:0), TG(10:0/10:0/14:1), TG(16:0/18:2/18:3), TG(18:2/18:2/18:3), TG(16:0/18:2/18:2), TG(16:0/18:1/18:2), TG(10:0/10:0/12:0), TG(18:2/18:2/18:2), TG(15:0/18:2/18:3), TG(18:1/18:2/18:2),DG(16:0/18:1), and TG(16:0/16:0/18:2) were significantly higher in the stem of the tolerant cultivar while TG(10:0/10:0/14:0), DG(18:3/18:3), TG(16:0/16:0/18:3), TG(16:0/18:3/18:3), TG(18:3/18:2/18:3), TG(10:0/10:0/10:0), TG(8:0/8:0/8:0), TG(18:1/18:1/18:1), TG(18:1/18:1/18:2) and TG(16:0/16:0/18:2) were significantly higher in the stem of the susceptible cultivar in response to infection (Fig. 5d). These results showed that there were significantly higher levels of TG and DG molecular species in root and stem of tolerant cultivar challenged with the pathogen compared to the stem of the susceptible cultivar following infection.

### Lipid biochemical network demonstrating from a system biology perspective how the tolerant and susceptible soybean cultivars respond to *P. sojae* infection

Lipid structural similarity networks were used to visualize changes in soybean root and stem lipids. For instance, the networks display three major clusters including top left (PSTs), top right (DGs and TGs containing saturated FAs), and bottom (a mixture of GPLs, DGs and TGs containing unsaturated FAs. CME 20:3 is the precursor for the biosynthesis of all the PSTs in the pathway presented, the level was significantly decrease resulting in downstream decrease in all unsaturated acylated hexocyl sitosterols. StE 18:3 had the biggest decrease in the ORC vs. ORI network of PST. In contrast, StE 18:3 increased several folds in CRC vs. CRI network, and it had the biggest increase. Generally, almost all the PSTs were decreased in the tolerant cultivar in response to infection. In the ORC vs ORI network, TG8:0/8:0/8:0, TG18:0/16:0/18:1, TG16:0/18:3/18:3, TG16:0/18:3/18:3 and TG16:0/16:0/18:3 are unique biomarkers differentiating the ORC vs. ORI while TG10:0/11:0/20:5 and DG18:0/18:0 were unique biomarkers differentiating CRC vs. CRI (Fig. 6). In OSC vs. OSI, StE 18:3 is a precursor for biosynthesis of all the PSTs, the level was significantly reduced leading upstream increase in all unsaturated acylated hexocyl sitosterols. AcHexSiE18:2 and AcHexSiE18:1 was increased several folds in CRC vs. CRI network. Similar to the root, almost all the PSTs in stem were reduced in the tolerant cultivar compared to the susceptible cultivar. In OSC vs. OSI, DG22:0/18:2 was the only unique biomarker differentiating OSC vs. OSI while in the CSC vs. CSI, TG12:0/12:0/12:0, TG16:0/16:0/18:2, TG10:0/10:0/14:1 and DG20:0/22:0 were unique biomarkers differentiating CSC vs. CSI (Fig. 7). In the ORI vs. CRI, TG10:0/10:0/10:0, TG15:0/22:0/22:5, DG 18:3/18:3 and DG16:0/18:3 were unique biomarkers differentiating ORI vs. CRI and TG10:0/10:0/14:0 and DG24:0/18:2 were unique biomarkers differentiating OSI vs. CSI (Fig. 8). Lipid species that changed only within one of these comparisons when considering all other comparisons (root and stem combined) are denoted with hashed outlines and may identify unique markers representative of the biological changes between these groups (Supplemental Table 1).

**Fig. 6.**
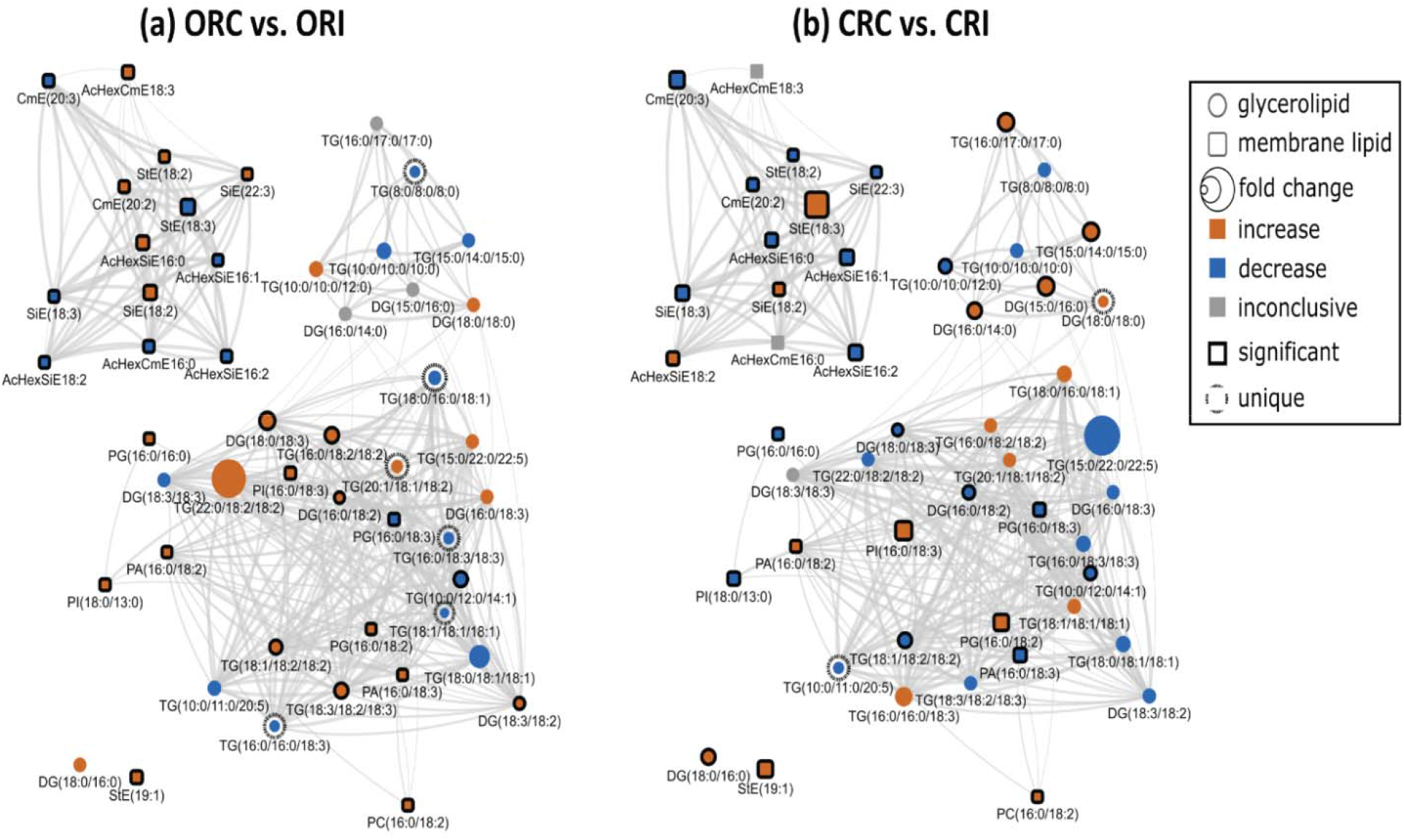
Lipid biochemical network displaying differences in storage and membrane lipids in the root of susceptible and resistant soybean cultivars inoculated with *P. sojae* relative to control plants. (**a**) Control susceptible soybean cultivar (ORC) versus inoculated (ORI); (**b**) control tolerant soybean cultivar (CRC) versus inoculated (CRI). The lipid biochemical network demonstrates fold differences in 22 root membrane lipid molecular species and 27 glycerolipid molecular species following inoculation with *P. sojae*. Lipid SMILES identifiers were used to calculate PubChem molecular fingerprints and structural similarities. Mapped networks, displaying significance of fold differences in lipids were calculated for all comparisons. Network visualizations display lipids connected based on structural Tanimoto similarity ≥ 0.8 (edge width: 0.8 to 1.0). Node size displays fold differences of means between comparisons and color shows the direction of change compared to control (orange: increased; blue: decreased; gray: inconclusive). Node shape displays lipid structural type (rounded square: membrane lipids; circle: glycerolipids). Lipids displaying significant differences between treatment groups (*p* ≤ 0.05) are denoted with black borders.

**Fig. 7.**
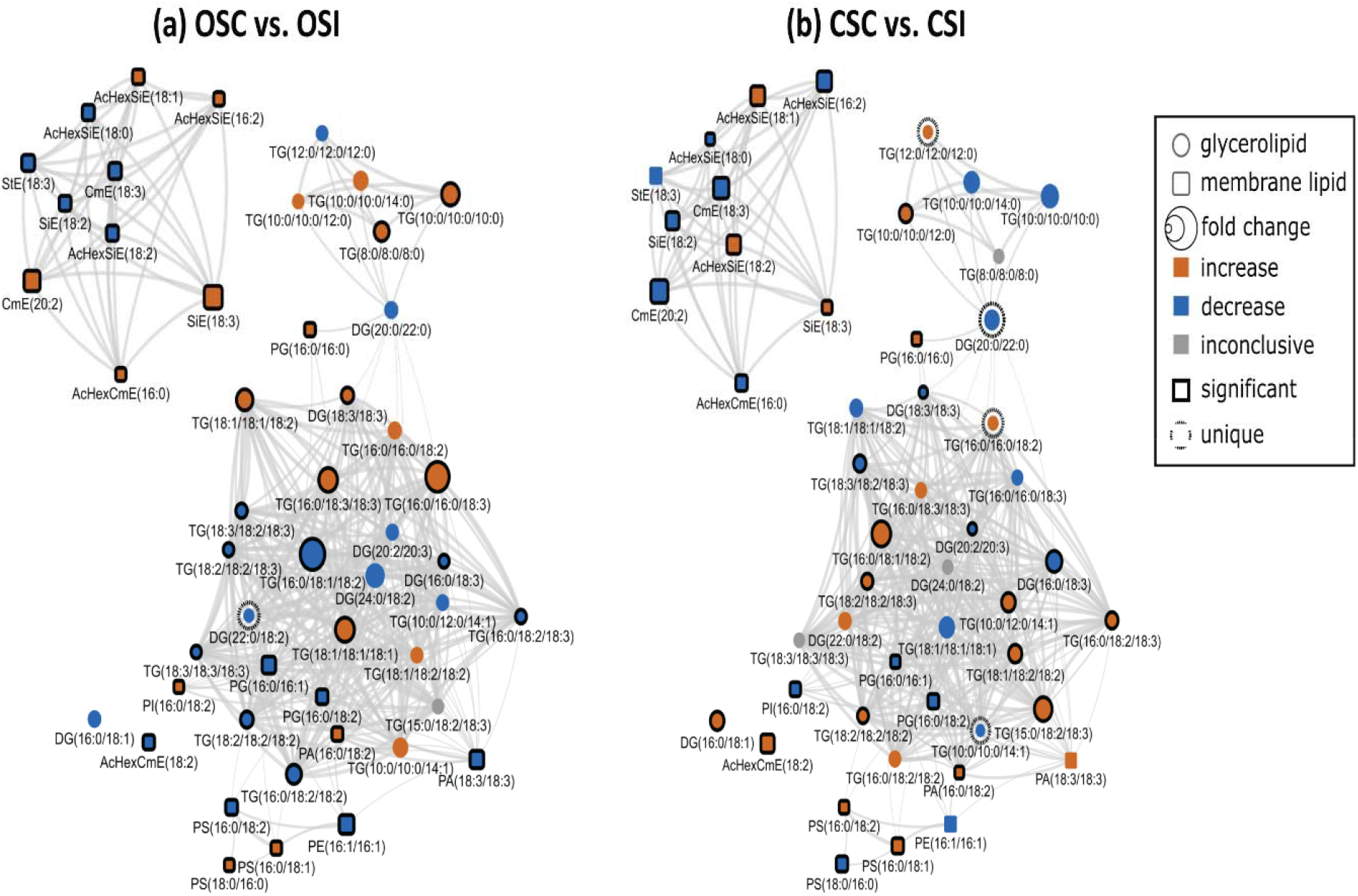
Lipid structural similarity network displaying differences in stem membrane lipids and glycerolipids in susceptible and resistant soybean cultivars inoculated with *P. sojae* relative to control plants. (**a**) Control susceptible soybean cultivar (OSC) versus inoculated (OSI); (**b**) control tolerant soybean cultivar (CSC) versus inoculated (CSI). The biochemical lipid network demonstrates fold differences in 21 stem membrane lipid molecular species and 28 glycerolipid molecular species following inoculation with *P. sojae*. Lipid SMILES identifiers were used to calculate PubChem molecular fingerprints and structural similarities. Mapped networks, displaying significance of fold differences in lipids were calculated for all comparisons. Network visualizations display lipids connected based on structural Tanimoto similarity ≥ 0.8 (edge width: 0.8 to 1.0). Node size displays fold differences of means between comparisons and color shows the direction of change compared to control (orange: increased; blue: decreased; gray: inconclusive). Node shape displays lipid structural type (rounded square: membrane lipids; circle: glycerolipids). Lipids displaying significant differences between treatment groups (*p* ≤ 0.05) are denoted with black borders.

**Fig. 8.**
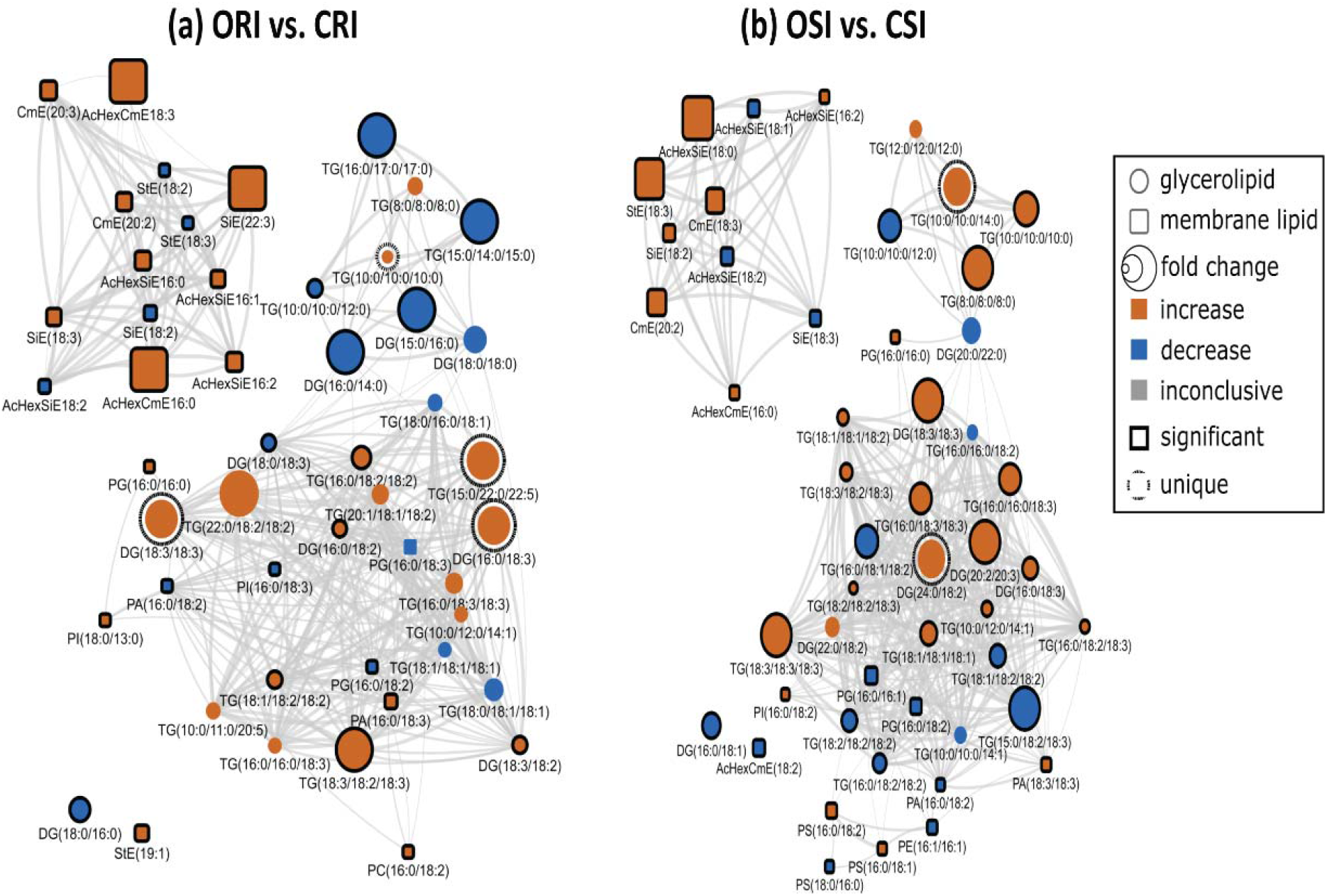
Lipid structural similarity network displaying differences in root and stem membrane lipids and glycerolipids in susceptible and resistant soybean cultivars inoculated with *P. sojae*. (**a**) Lipids from inoculated root tissue of susceptible (ORI) versus tolerant (CRI) soybean cultivars inoculated with *P. sojae*; and (**b**) Lipids from inoculated stem tissue of susceptible (OSI) versus tolerant (CSI) soybean cultivars inoculated with *P. sojae*. The biochemical lipid network demonstrates fold changes in 22 root membrane lipid molecular species and 27 glycerolipid molecular species, and 21 stem membrane lipid molecular species and 28 glycerolipid molecular species following inoculation with *P. sojae*. Lipid SMILES identifiers were used to calculate PubChem molecular fingerprints and structural similarities. Mapped networks, displaying significance of fold differences in lipids were calculated for all comparisons. Network visualizations display lipids connected based on structural Tanimoto similarity ≥ 0.8 (edge width: 0.8 to 1.0). Node size displays fold differences of means between comparisons and color shows the direction of change compared to control (orange: increased; blue: decreased; gray: inconclusive). Node shape displays lipid structural type (rounded square: membrane lipids; circle: glycerolipids). Lipids displaying significant differences between treatment groups (*p* ≤ 0.05) are denoted with black borders.

## Discussion

As essential components of cellular membranes, lipids are involved in various physiological roles including as structural components of cellular membranes, cell signaling, storage of energy, and membrane trafficking. In plants, alterations in lipid composition have been reported in response to pathogenic stress conditions (Nurul Islam et al. 2012). Biotic stress have been reported to profoundly alter the lipidome in plants (Kim et al. 2013). Additionally, Ferrer et al. (2017) demonstrated that alterations in the relative composition of PSTs in cellular membranes affect their biophysical properties and hence their physiological functions. The results we describe here indicate significant alterations in lipid metabolism in both a resistant and a susceptible soybean cultivar in response to *P. sojae* infection. Specifically, in the pathogen-treated plants, we observed significantly higher levels of major GPLs and GLs (DGs and TGs) in the tolerant cultivar, whereas StEs and CmEs were found to be higher in quantity in the susceptible cultivar. More interestingly, these classes of lipids varied in a similar manner in the root and stem of each cultivar in response to pathogen infection, which is in line with the literature (Kim et al. 2013, Naguib 2019, Shah 2005). For example, similar trends were observed for the lipidome of eggplants (*Solanum melongena*) resistant to Fusarium wilt infection (Naguib 2019), demonstrating the significant difference in the levels of lipid profiling of the susceptible and tolerant eggplants to Fusarium disease and this ensured the essential roles of the lipids in resistance strategy against infection (Naguib 2019). The increased lipid levels in tolerant cultivars serve as energy stores and provide a buffer to stress; the stored lipids could act as additional energy that keeps the plants from shifting to proteolysis and then cell death (Shah 2005).

The biosynthesis and lipid composition of cellular membranes play an essential role in the physiological functioning of plants (Reszczyńska and Hanaka 2020). During growth, plants adapt to adverse stress conditions through the remodelling of lipid membranes resulting from alterations in the fatty acid content and, consequently, the biosynthesis of lipids (Reszczyńska and Hanaka 2020). Several studies have demonstrated that high levels of lipid remodeling in plant membrane lipids under different adverse conditions result in resistance to environmental stressors (Reszczyńska and Hanaka 2020).

Our results clearly show that there are differences in both membrane and storage lipid metabolism in resistant and susceptible soybean cultivars in response to *P. sojae* infection. For instance, we observed higher levels of 18:2 and 18:3 fatty acyl-enriched phospholipid and sterol molecular species in the membrane lipids of the root and stem from the tolerant cultivar when challenged with the pathogen, in contrast to lower C18:2 and C18:3-enriched molecular species in tissues from the susceptible cultivar (Fig. 2, 3). In contrast, the StEs were significantly higher in the root and stem from the susceptible cultivar challenged with *P. sojae* infection but were significantly lower in the susceptible control plants and in the tolerant cultivar under both treatment conditions (Table 1, 2; Fig. 2, 3). This is in agreement with a recent study which demonstrated the role of sterols in disease resistance (Kopischke et al. 2013). Stigmasterol ester was identified as a factor of susceptibility in *Arabidopsis*, as inhibition of its biosynthesis resulted in increased resistance to *Pseudomonas syringae* (Griebel and Zeier 2010, Kopischke et al. 2013). Another report indicated that C22 desaturation of the main phytosterol, β-sitosterol, in *Arabidopsis* through the enzyme CYP710A1, and the associated stigmasterol accumulation, are important metabolic activities in *P. syringae*-inoculated leaves of *Arabidopsis* that can increase susceptibility (Griebel and Zeier 2010). The formation of stigmasterol in leaves is induced by recognition of bacterial pathogen-associated molecular patterns and synthesis of reactive oxygen species, but is independent of the jasmonic acid, salicylic acid or ethylene-associated signalling pathways (Griebel and Zeier 2010). Through analysis of mutants and application of exogenous sterol, it was demonstrated that an increase in the ratio of stigmasterol to β-sterol in leaves reduces specific defence responses in *Arabidopsis*, and consequently makes the plants more susceptible to *P. syringae* (Adigun et al. 2020, Wang et al. 2012). These were in line with the results obtained in this study, and these modes of action may account for the higher resistance of the tolerant cultivar to pathogen infection.

Pathogenic fungi can secrete various extracellular enzymes that are involved in pathogenicity (Subramoni et al. 2010). For example, secreted lipases from fungal pathogens are involved in the penetration of plant barriers such as the wax cuticle. Likewise, internal fungal lipases are capable of degrading storage lipids and/or signaling via the release of secondary messengers. The significant decrease in the TG molecular species in the soybean susceptible cultivar could be as a result of increased lipase activity during infection. Lipases hydrolyze carboxyl esters in TGs and liberate FAs and glycerol (Watt and Steinberg 2008). This is in agreement with the fact that lipases appear to function as virulence factors in plant pathogens. Interestingly, in this study, the tolerant cultivar demonstrated significantly higher DG levels in response to pathogen infection, but there was no observed difference in TG levels. This is in agreement with the fact that DGs are primarily derived either from TGs through TG lipases or PAs by phospholipase activity (Bates and Browse 2012). However, it has also been reported that DG levels in a tolerant eggplant cultivar can be generated by the activity of phospholipase on PAs, and not only the activity of TG lipases (Naguib 2019).

The lipid biochemical network demonstrated significant alterations in lipid metabolism in both cultivars in response to *P. sojae* infection. The head group and FA composition of complex lipids are a useful proxy for localization and biological function (Casares et al. 2019). Networks display increased density in connectivity between biochemically related groups of lipids and the lipid biosynthesis metabolism pathway in the tolerant soybean cultivar as defense response to pathogen inversion. Generally, there is dearth of information on the role of lipid metabolism in determining either incompatible or compatible interactions in the soybean-*P sojae* pathosystem during host-pathogen interaction. The unique biomarkers between the susceptible and tolerant cultivars including the production of DG molecular species, which was well pronounced in tolerant cultivar than susceptible (Figs. 6, 7 and 8). Studies have demonstrated that signaling enzymes, diacylglycerol kinases (DGKs) play important roles in response to biotic stress by phosphorylating DG to synthesis PA (Fig. 9) and both PA and DG are lipid mediators during physiological process (Yuan et al. 2019). Our findings from this study demonstrate that lipid metabolism and signalling possibly involving DG could play a significant role in pathogen resistance in the tolerant soybean cultivar. Also, DG signally related to TG hydrolysis which was differentially demonstrated between susceptible and tolerant soybean cultivars when challenged with pathogen (Figs 6, 7 and 8). Study has demonstrated that TG is accumulated in plant tissues due to TG turnover, as a result of disruption of SUGAR-DEPENDENT1, a cytosolic lipase accountable for TG hydrolysis in lipid droplets into free FAs and DG and consequently enhance TG accumulation in plant tissues (Kelly et al. 2013). Fan et al (2017) demonstrated that TG accumulation plays important role, thus buffering homeostasis of lipid and protecting plant cells against lipotoxic death as a results FA overload and can be as a remodeling of robust membrane in response to stresses. Phytosterols also known to play important role in plant innate immunity against pathogen attack (Wang et al. 2012).

**Fig. 9.**
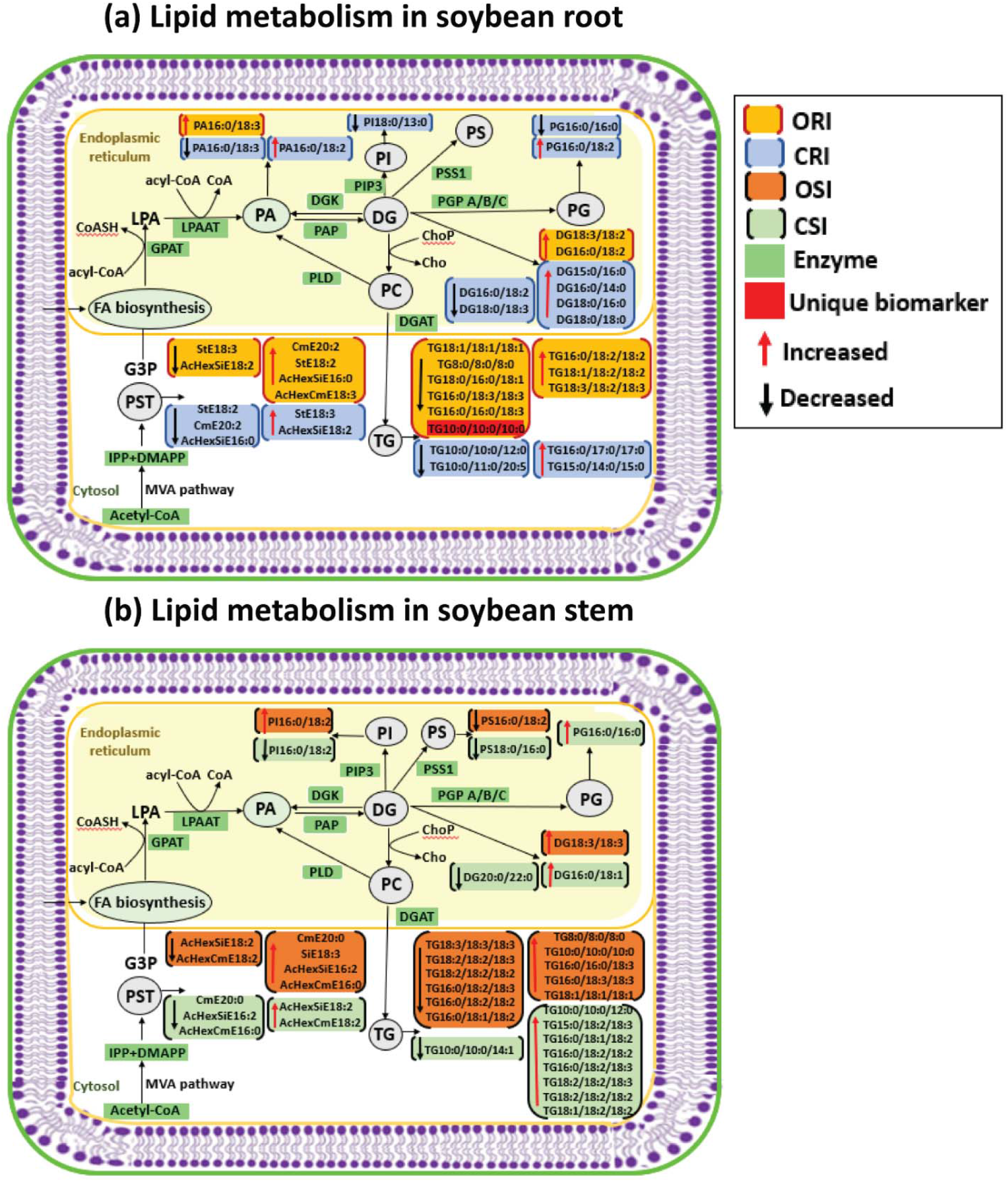
Proposed lipid metabolism pathways suggesting the mechanism that maybe associated with the altered lipidome and disease tolerance or susceptibility in soybean cultivars (OX760-6 and Conrad) following inoculation with *P. sojae*. (a) The most significantly altered root lipids in soybean cultivars (OX760 and Conrad) in response to colonization and infection with P.sojae; and (b) The most significantly altered stem lipids in soybean cultivars (OX760 and Conrad) in response to *P. sojae* colonization and infection. In the Kennedy pathway fatty acyl-CoA and coenzyme A begins with the sequential acylation of GPATs and LPAATs utilizing fatty acyl-CoA to biosynthesis the central precursor PA through which other downstream GPLs are produced. GLPs are produced through hydrolysis of the phosphate group in PA, and this PA then dephosphorylated through PAP to generate DG. The DG acts as a precursor for biosynthesis of TG via DGAT or PDAT transferring the sn-2 fatty acyl group from GPLs to DG, producing TG. Biosynthesis of IPP and DMAPP through mevalonate (MVA) pathway, and they act as precursors for phytosterol synthesis. The altered lipidome observed in this study suggest DG and PA mediated lipid signalling impacting phytosterol anabolism appears to be the strategy used by tolerant soybean cultivars to successfully limit infection and colonization by *P*.*sojae*. The following molecular species are suggested as unique lipid biomarkers in the ORI vs CRI and CSI vs OSI networks that could potentially discriminate tolerance interations in the soybean-P.sojae pathosystem: TG(15:0/22:0/22:5), TG(10:0/10:0/10:0), TG(10:0/10:0/14:0), DG(18:3/18:3), DG(16:0/18:3) and DG(24:0/18:2). PLD = phospholipase D, DGK = diacylglycerol kinase, LPAAT = lysophosphatidic acid acylteransferase, PAP = phosphatic acid phosphatase, G3P = glycerol-3-phosphate, DGAT = diacylglycerol acyltranferase, GPAT = Glycerol-3-phosphate acyltransferase, PDAT = phospholipid:diacylglycerol acyltransferases, PSS1 = phosphatidylserine synthase-1, PGP = glycerol-3-phosphate phosphatase, PAP = phosphatidic acid phosphatase, IPP = isopentenyl pyrophosphate, DMAPP = dimethylallyl pyrophosphate, MVA = mevalonic acid, PIP3 =1-phosphatidylinositol-4-phosphate 5-kinase, CoASH = coenzyme A, Chop = cholinephosphotransferase and cho = choline. ORI = root of susceptible inoculated, CRI = root of tolerant inoculated, OSI = stem of susceptible inoculated, CSI = stem of tolerant inoculated, GPLs = glycerophospholipids, GLs = glycerolipids, LPA = lysophosphatidic, PA = phosphatidic acid, PC = phosphatidylcholine, PG = phosphatidyl glycerol, PI = phosphatidylinositol, PS = phosphatidylserine, DG = diacylglycerol, TG = triacylglycerol and PST = phytosterols.

Lipid biosynthesis in soybean cultivars follow common routes where FAs are generated from plastid, transported to the endoplasmic reticulum (ED) (Kim 2020), which starts with the addition of fatty acyl-CoA leading to biosynthesis of lysophosphatidic acid (LPA) and the reaction is catalyzed by glycerol phosphate acyltransferase (GPAT) and is a rate limiting-step for PA biosynthesis. In ED, PA biosynthesis occurs by addition of fatty acyl-CoA to LPA via lysophosphatidic acid acyltransferase (LPAAT) to form central precursor PA by which several GPLs are synthesized (Fig. 9). The first step in GPLs biosynthesis involves the hydrolysis of the phosphate group from PA to generate DG by phosphatidic acid phosphatase (PAP). The resulting DG is later phosphorylated by DGKs to PA, which is subsequently reused in biosynthesis of GPLs. Also, DG acts as a precursor for biosynthesis of primary form of storage energy, TG (Kelly and Jacobs). The isopentenyl diphosphate (IPP) and dimethylallyl pyrophosphate (DMAPP) generated via cytosolic mevalonate (MVA) pathway are primarily used for the biosynthesis of phytosterols (Lohr et al. 2012, Vriese et al. 2019). Our results demonstrate novel information about pathogen-stress responses in the root and stem of both soybean cultivars, which can be put within the broad context of plant lipid metabolism. The metabolic pathway of relative abundance of GPL, PST and GL biosynthesized in the root and stem of the susceptible and tolerant soybean cultivars when challenged with *P. sojae* are demonstrated in Fig. 9. These lipid classes could be used as biomarkers for disease resistance or susceptibility by soybean cultivars. Based on our understanding, this is the first report of lipid alteration in soybean root and stem in response to *P. sojae* infection.

To conclude, our results demonstrate promise for a novel mechanism to engineer soybean cultivars for wide spectrum disease susceptibility or resistance by manipulating plant lipid levels. Both soybean cultivars altered lipid biosynthesis upon infection by *P. sojae*. Induced accumulation of stigmasterol, and total increase in the ratio of stigmasterol to β-sitosterol in the susceptible soybean cultivar favoured pathogen multiplication and then improved disease susceptibility whereas induced accumulation and overall increase in GPLs (PA and PG) and GLs (DG and TG) in tolerant soybean cultivar enhanced plant immunity against pathogen.

Glycerophospholipids strengthen the cellular membrane and protect plant cells from various infections while DGs mainly act as signalling molecules during response to various environmental stresses. The altered lipidome observed in this study suggest DG and PA mediated lipid signalling impacting phytosterol anabolism appears to be the strategy used by tolerant soybean cultivars to successfully limit infection and colonization by *P*.*sojae*. The following molecular species are suggested as unique lipid biomarkers in the ORI vs CRI and CSI vs OSI networks that could potentially discriminate tolerance interactions in the soybean-P.sojae pathosystem: TG(15:0/22:0/22:5), TG(10:0/10:0/10:0), TG(10:0/10:0/14:0), DG(18:3/18:3), DG(16:0/18:3) and DG(24:0/18:2). To understand the exact roles of these plant lipids in membrane permeability and as signaling molecules warrant further studies.

## Materials and methods

### Plant growth and inoculation method

A virulent strain of *P. sojae* race 2 (strain P6497) obtained from the London Research and Development Center, Agriculture and Agri-Food Canada (AAFC-LRDC; London, ON, Canada) was used as inoculum. The oomycete was cultured and maintained aseptically for 8 days on 26% V8-juice agar (8400 mg agar, 1600 mg CaCO_3_, 156 mL V8-juice [Campbell Soup Company, Toronto, ON, Canada], and 440 mL of distilled water). Seeds of soybean cultivars Conrad (*P. sojae-*tolerant) and OX760-6 (*P. sojae-*susceptible) were obtained from AAFC-LRDC (London, ON, Canada). The seeds were surface disinfected for 5 min using 0.5% sodium hypochlorite (Commercial Javex Bleach; Clorox Co., Brampton, Ontario, Canada) and rinsed with distilled water several times. The seeds were then soaked for 12 h in distilled water before seeding. The bottom of a sterilized empty paper drink cup was used to cut agar disks consisting of cultures of *P. sojae* P6497 which were then fitted into the bottom of wax-paper cups with a top diameter of 8.5 cm and 15.0 cm deep (Merchants Paper Company, Windsor, ON, Canada) and overlaid with medium-grade vermiculite. Drainage holes were created in the bottom of the cups. The imbibed seeds were planted in the medium-grade vermiculite. Six soybean seedlings from each cultivar were inoculated with *P. sojae* in a cup and another six from each cultivar were mock-inoculated (sterile V8-juice agar disks without a *P. sojae* culture) in a cup as the control. The plants were then grown for 10 days. The plant growth experiment was performed in a growth chamber (Biochambers MB, Canada) at Grenfell Campus, Memorial University of Newfoundland, under controlled growth conditions of 16 h light at 25°C and 8 h dark at 20°C, and relative humidity of 60%. Seedlings were watered daily 4 days after seeding with one-quarter-strength Knop’s solution (Thomas et al. 2007). The whole seedlings were collected 10 days after growth and stored at -80°C until further analysis.

## Method of lipid extraction

Soybean seedlings prepared as above were incubated in boiling isopropanol for 10 min. Lipid extraction was conducted by weighing 100 mg each of root and stem from each sample type, and 1 mL MeOH containing 0.01% butylated hydroxytoluene was added to each sample. Four replications of each combination of treatment (inoculated or control), cultivar (susceptible or tolerant), and tissue (root or stem) combination were performed. The tissues were then homogenized using a probe tissue homogenizer until completely dissolved. Following homogenization, 800 µL water and 1000 µL chloroform were added along with PC 14:0/14:0 as internal standard. Each sample was thoroughly vortexed and centrifuged at 3000 rpm for 15 min at room temperature. The organic layers were transferred to pre-weighed 4 mL glass vials with PTFE-lined caps (VWR, Mississauga, Canada). The samples were then dried under a gentle stream of nitrogen and the sample vials reweighed to determine the quantity of recovered lipids. The recovered lipids from each sample were re-suspended in 1000 µL solvent (2:1 v/v chloroform: methanol) and stored at -20°C until lipid analysis using ultra high-performance liquid chromatography coupled to heated electrospray ionization high resolution accurate mass tandem mass spectrometry (UHPLC-C30RP-HESI-HRAM-MS/MS).

### Lipid analysis using UHPLC-C30RP-HESI-HRAM-MS/MS

The method of lipid analysis was as described previously (Nadeem et al. 2020). Lipids extracted from the soybean roots and stems were separated using an Accucore C30 reverse phase (C30RP) column (150 × 2 mm I.D., particle size: 2.6 µm, pore diameter: 150 Å; ThermoFisher Scientific, ON, Canada) applying the following solvent system: Solvent A (40: 60 v/v H_2_O and acetonitrile), and Solvent B (1:10: 90 v/v/v water: acetonitrile: isopropanol). Both solvents A and B consisting of 0.1% formic acid and 10 mM ammonium formate. The conditions for the separation using UHPLC-C30RP were as follows: oven temperature of 30°C, flow rate of 0.2 mL/min, and 10 µL of the lipid mixture suspended in 1: 2 v/v methanol: chloroform was injected into the instrument. The system gradient used for the separation of lipid classes and molecular species were: 30% solvent B for 3 min; solvent B increased over 5 min to 43%, then increased in 1 min to 50% B and to 90% B over 9 min; and from 90% to 99% B over 8 min; and finally maintained at 99% B for 4 min. The column was re-equilibrated to 70% solvent A for 5 min to re-establish the starting conditions before injection of each new sample. Lipid analyses were performed using a Q-Exactive Orbitrap high-resolution accurate mass tandem mass spectrometer (Thermo-Scientific, Berkeley, CA, USA) coupled with an automated Dionex Ulti-Mate 3000 UHPLC system controlled by Chromeleon 6.8 SR13 (Dionex Corporation, Part of Thermo Fisher Scientific) software. Full-scan HESI-MS and MS/MS acquisitions were performed in positive mode of the Q-Exactive Orbitrap mass spectrometer. The following parameters were used for the Orbitrap mass spectrometry techniques: auxiliary gas of 2; sheath gas of 40; capillary temperature of 300°C; ion spray voltage of 3.2 kV; S-lens RF of 30 V; full-scan mode at a resolution of 70,000 m/z; mass range of 200–2000 m/z; top-20 data dependent MS/MS acquisitions at a resolution of 35,000 m/z; and injection time of 35 min; automatic gain control target of 5e5; isolation window of 1 m/z; collision energy of 35 (arbitrary unit). The external calibration of instrument was performed to 1 ppm using ESI positive and negative calibration solutions (Thermo Scientific, Berkeley CA, USA). Mixtures of lipid standards were used to optimize tune parameters (Avanti Polar Lipids, Alabaster, AL, USA) in both positive and negative ion modes. Identification and semi-quantification of the classes of lipids and lipid molecular species present in the root and stem of both soybean cultivars (OX760-6 and Conrad) were performed using LipidSearch version 4.1 (Mitsui Knowledge Industry, Tokyo, Japan) and the parameters adopted for identification in LipidSearch were: target database of Q-Exactive; product tolerance of 5 ppm; precursor tolerance of 5 ppm; Quan m/z tolerance of ±5 ppm; product ion threshold of 5%; m-score threshold of 2; Quan retention time range of ±1 min; use of all isomer filter; ID quality filters A, B, and C; and [M+NH_4_]^+^ adduct ions for positive ion mode. Following identification, the observed lipid classes and lipid molecular species were merged and aligned according to the parameters established in our previous report (Pham et al. 2019).

### Lipid biochemical network mapping

To better understand how soybean cultivars that are tolerant and susceptible to *P. sojae* modulate their membrane lipid metabolism as part of the plant defense response strategy during infection and colonization, lipids that changed significantly between treatments were visualized within lipid structural similarity and implied activity networks. Lipid SMILES identifiers obtained from lipid map were used to calculate PubChem molecular fingerprints describing lipids’ sub structures (Guha 2007). Connections between lipids were defined based on Tanimoto similarity ≥ 0.8 between fingerprints. Significance of fold changes in lipid expression levels were mapped to network node attributes and displayed using Cytoscape (Grapov et al. 2015, Shannon et al. 2003). Node size was used to represent fold changes of means between treatments, and colors indicated the direction of change compared to control (orange = increased; blue = decreased; gray = inconclusive) in the lipid network map generated. Node shape was used to indicate lipid structural type (rounded square= membrane lipids; circle = neutral lipids). Lipids displaying significant differences between treatment groups (p ≤ 0.05) were denoted with black borders.

### Statistical analysis

To determine the effects of pathogen infection on lipid composition of the root and stem of susceptible and tolerant cultivars, multivariate analyses including partial least square discriminant analysis (PLS-DA), and heat map were performed to group the treatments based on similarity. Analysis of variance (ANOVA) was next performed to determine whether the groups were significantly different between treatments using XLSTAT (2017 Premium edition, Addinsoft, Paris, France). Where significant differences were observed, the means were compared with Fisher’s Least Significant Difference (LSD), α = 0.05. Figures were prepared with SigmaPlot 13.0 (Systat Software Inc., San Jose, CA).

## Author contribution

OAA performed experimental analysis. OAA, THP, DP, MN worked on data curation. OOA drafted the manuscript. RM, LEJ, MC, LG planned and designed the research. All co-authors reviewed and provided input to the final manuscript.

## Acknowledgements

We acknowledge Dr. Mark Gijzen (Agriculture and Agri-Food, Canada, London Ontario) for providing soybean seeds and Dr. Tao Yuan proper maintenance of laboratory instruments.

